# Simultaneous mapping of RNA secondary, tertiary, and quaternary structure in living cells by multi-site DMS probing

**DOI:** 10.1101/2024.11.27.625770

**Authors:** Irfana Saleem, Thomas Miller, Lucas Kearns, David Mitchell, Ritwika Bose, Chase A. Weidmann, Anthony M. Mustoe

**Affiliations:** Therapeutic Innovation Center (THINC), Verna and Marrs McLean Department of Biochemistry and Molecular Pharmacology, Baylor College of Medicine, Houston, TX; Department of Biological Chemistry, Center for RNA Biomedicine, Rogel Cancer Center, University of Michigan Medical School, Ann Arbor, MI; Department of Molecular and Human Genetics, Baylor College of Medicine, Houston, TX

## Abstract

Identifying tertiary structures and protein binding sites on RNA molecules remains a key challenge in RNA biology. We report a new chemical probing strategy termed multi-site DMS-MaP (msDMS-MaP) that exploits base tautomerization during mutational profiling reverse-transcription to measure typically invisible DMS-induced modifications at the N7 position of G. These N7-G modifications are now resolved concurrently with conventional N1 and N3 modifications with only minor modifications to established protocols, providing a multi-dimensional, single-molecule readout of RNA structure. We show that N7-G reactivity specifically reports on RNA tertiary and quaternary structure, enabling identification of diverse, functionally significant motifs such as cooperatively folding tertiary domains and protein binding sites in living cells. We apply msDMS-MaP to obtain a map of the quaternary structural ensemble of the 7SK non-coding snRNP, revealing unique protein binding sites across three 7SK structural isoforms. msDMS-MaP represents a broadly applicable strategy for enhanced RNA functional motif discovery and characterization.

## INTRODUCTION

RNA molecules function via complex secondary and tertiary structures that specifically interact with proteins, small molecule ligands, and other cellular effectors^1^. In many cases, these RNA structures and quaternary complexes are dynamic, cycling between different RNA conformations and molecular interactions that support distinct functional outcomes^2–4^. Defining RNA tertiary and quaternary structures and their dynamics, particularly in living cells, is a major challenge in RNA biochemistry, representing a critical bottleneck for understanding biological mechanisms and for efforts to therapeutically target RNA.

Current RNA biochemical characterization strategies provide information about either secondary structure, tertiary structure, or protein or other ligand binding sites, but typically not concurrently^5^. Dimethyl sulfate (DMS) and 2′-hydroxyl acylation (SHAPE) reagents are widely used to measure RNA secondary structure by selectively modifying single-stranded nucleotides followed by sequencing-based readout such as mutational profiling (MaP)^5–8^. In principle, these experiments also encode information about RNA tertiary structures and ligand binding sites, which can protect RNA nucleotides from modification^9–11^. However, deconvoluting whether nucleotides are unreactive due to rare tertiary and quaternary interactions versus ubiquitous secondary structures is challenging, requiring multiple comparative experiments. Other methods that can directly map RNA tertiary and quaternary structures provide limited to no information on secondary structure^12–17^. Thus, holistic RNA structural characterization typically requires multiple laborious and expensive experiments. More significantly, conformationally dynamic RNAs often undergo coordinated changes in secondary and higher-order structure^2,3^, and resolving such coordinated changes is especially difficult with current methods.

Here, we introduce a new DMS chemical probing strategy that enables concise, concurrent measurement of RNA secondary and higher-order structural ensembles in living cells. Standard DMS probing experiments focus on detecting DMS-induced methylations at purine N1 and pyrimidine N3 positions, which are readily mapped by reverse-transcriptases and provide sensitive measures of secondary structure^7,8,18^. However, the most prevalent DMS methylation is at the N7 position on the Hoogsteen face of guanine (G) nucleobases (Fig 1A)^19–21^. N7-G reactivity provides information about RNA higher-order structure^19,22^, but measuring N7-G modifications has historically required harsh, multi-step biochemical processing^19,22–26^, and the value of N7-G reactivity for RNA structure characterization is underexplored. We developed a simple reverse-transcription strategy that exploits tautomerism of N7-modifed bases to enable high-efficiency, concurrent detection of N7-G and N1 and N3 modifications, which we term multi-site DMS probing readout by mutational profiling (msDMS-MaP). We demonstrate that N7-G reactivity provides a highly specific measure of RNA higher-order structure, enabling both high-throughput discovery and detailed characterization of diverse tertiary structures and protein binding sites across RNA systems. We then apply msDMS-MaP to map the quaternary structural ensemble of the human 7SK ribonucleoprotein (RNP) in living cells. By providing additional tertiary and quaternary detail to the robust DMS-MaP strategy, msDMS-MaP promises to broadly accelerate RNA functional motif discovery and mechanistic analysis.

**Figure 1:**
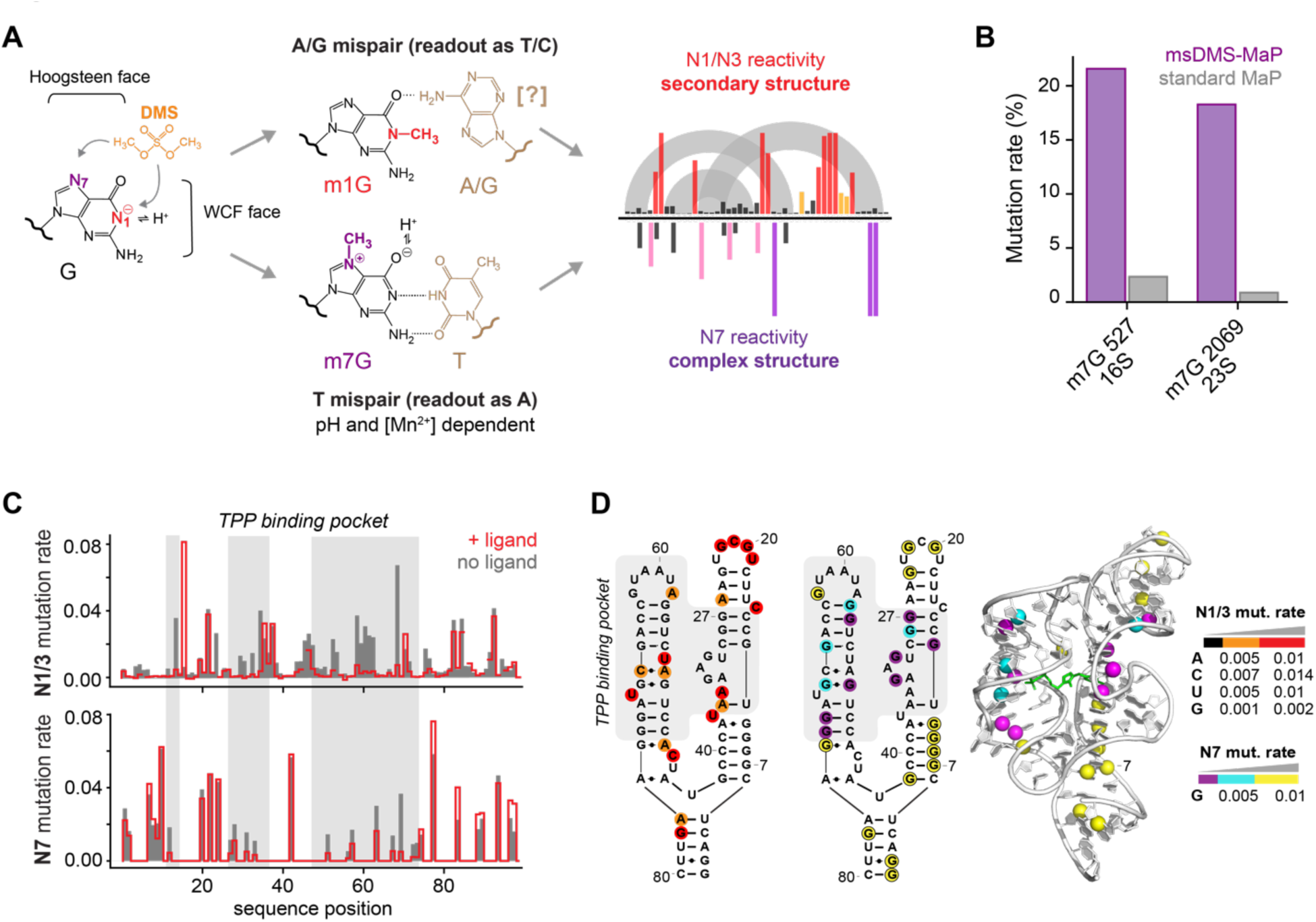
Tautomer stabilization permits simultaneous measurement of multiple DMS modifications by mutational profiling reverse transcription. (A) Schematic and proposed mechanism by which MarathonRT decodes DMS-induced methylations at N1- and N7-G positions. N1-G modifications at the Watson-Crick-Franklin (WCF) face induce A/G mispairs (read as G→U/C mutations^18^), possibly via a syn pairing mechanism. N7-G modifications at the Hoogsteen face stabilize the enol and enolate tautomers of guanine and induce T mispairs (read as G→A mutations). (B) Decoding efficiency of natural N7-G modifications in *E. coli* ribosomal RNA by msDMS-MaP and standard MaP protocols. (C) Background subtracted N1/3 mutation rates (top) and background subtracted N7 mutation rates (bottom) for the *E. coli* TPP riboswitch with and without ligand. Regions that are protected upon adding ligand are highlighted gray. (D) Left: secondary structure of the TPP riboswitch with highly, moderately, and lowly reactive N1/3 positions colored in red, orange, black, respectively. Middle: secondary structure of the TPP riboswitch with highly, moderately, and lowly reactive N7-G positions colored in yellow, cyan, and purple, respectively. Right: N7-G reactivity data superimposed on the TPP three-dimensional structure PDB 2GDI^32^. Reactivity rate cutoffs for each nucleotide are shown in the key at right.

## RESULTS

### Tautomeric stabilization permits reverse transcriptase decoding of m7G RNA modifications

We previously reported that m7G RNA modifications, while conventionally invisible to reverse-transcriptase enzymes (RTs), are inefficiently decoded by MarathonRT as C→T substitutions in cDNA^18^. These substitutions are subsequently resolved as a G→A mutational signature during sequencing. We hypothesized that this mutational signature arises from the increased susceptibility of m7G to form enolate and enol tautomers within the RT active site, facilitating canonical-like m7G·T pairing during RT decoding (Fig. 1A)^20,27,28^. Because m7G tautomerization is pH dependent^29^, we reasoned that we could selectively stabilize mutagenic m7G tautomers and increase the observed mutation rate by modulating the pH of the RT reaction. Indeed, we observed a strong pH-dependent and Mn^2+^-dependent increase in mutation rates at natural m7G bases with minimal changes in decoding fidelity at neighboring unmodified positions (Fig. 1B; S1A-D). Our improved RT buffer enabled detection of m7G modifications with ∼20% efficiency, a ∼15-fold improvement over standard SSII MaP reverse-transcription^18^ and comparable to the efficiency of first-generation depurination-based m7G-measurement strategies^23,26^. Unlike depurination strategies, tautomeric stabilization requires no chemical treatments prior to RT and thus is ideal for preserving RNA integrity and chemically labile modifications on the RNA.

We exploited this tautomer-stabilizing RT approach to develop a new DMS chemical probing strategy to concurrently interrogate RNA secondary and tertiary structure in a single experiment, which we term multi-site DMS probing readout by mutational profiling, or msDMS-MaP. Optimized DMS reaction conditions are used to modify the N7 position on the Hoogsteen face of G nucleobases and the N1 and N3 positions of the Watson-Crick-Franklin (WCF) faces of all four nucleobases, including U and G^18,30^. Importantly, N7-G modifications on the Hoogsteen edge produce distinct G→A mutational signatures during MaP RT, while N1-G modifications at the WCF face produce G→T/C mutational signatures (Fig. 1A)^18^. As in standard DMS-MaP, N1-A, N3-C, and N3-U modifications are also resolved as random mismatches at their respective nucleotide positions.

As an initial assessment, we collected msDMS-MaP data on the *E. coli* TPP riboswitch aptamer domain folded in solution with or without ligand^31,32^. N1/N3 modification rates measured with our msDMS-MaP protocol were highly correlated with modification rates obtained using standard MaP-RT (R>0.92; Fig. S1E) and conform to the known RNA secondary structure (Fig. 1C, D). In contrast, N7-G modification rates predominantly report on the riboswitch tertiary structure (Fig. 1C, D). For example, nucleotides G7-G10 are highly modified at the N7 position but unreactive at the N1 position, consistent with these nucleotides being base-paired but not involved in higher-order structures. By comparison, guanines within the ligand pocket are protected from N7 modification regardless of whether they are base-paired (G27, G28) or unpaired (G31, G33) (Fig. 1C, D). N7-G reactivity increases significantly in the absence of TPP ligand (Fig. 1C), confirming that N7-G reactivity depends on tertiary folding. Both N1/3 and N7-G reactivity profiles were highly reproducible across independent replicates (R>0.99; Fig. S1F). Together, these experiments support that msDMS-MaP accurately measures DMS modifications at both N1/N3 and N7 positions, providing complementary information on secondary and tertiary structure, respectively.

### N7-G protections identify complex folds and protein binding sites in the ribosome

To evaluate our msDMS-MaP strategy on a broader variety of RNA conformations, we collected msDMS-MaP data on the *E. coli* 16S and 23S ribosomal RNAs (rRNAs) under varied folding conditions: on fully assembled ribosomes in living *E. coli* cells; protein-free rRNA extracted from cells and folded in Mg^2+^–containing and Mg^2+^–free buffers to vary the degree of tertiary assembly; and urea-denatured rRNA extracted from cells (Fig. 2A). Measured N1/N3 and N7-G reactivities were highly replicable across independent experiments (R>0.97; Fig S2A, B). N1/N3 reactivities were also highly correlated with standard DMS-MaP measurements and equally predictive of secondary structure at all four nucleotides (Fig. S2C-E), confirming that our msDMS-MaP protocol supports high-fidelity N1/N3 measurement. N7-G reactivity was largely orthogonal to N1-G reactivity (R=0.33; Fig. S2F) and varied dramatically across conditions (Fig. 2B). Under denatured conditions, N7-G positions demonstrate a broad range of reactivity rates ranging from 0.01 to 0.15, with median reactivity of 0.05 (Fig. 2B). By comparison, most rRNA N7-G positions are unreactive in cells (rate < 0.01), consistent with tertiary structures and bound proteins providing extensive protection of N7-G sites. Protein-free rRNAs demonstrate intermediate levels of N7-G reactivity on average, indicating that WCF pairing moderately protects N7-G positions from DMS modification compared to fully single-stranded G nucleotides. Protein-free rRNAs also feature some lowly reactive N7-G sites, consistent with protection from self-folding tertiary structures.

**Figure 2:**
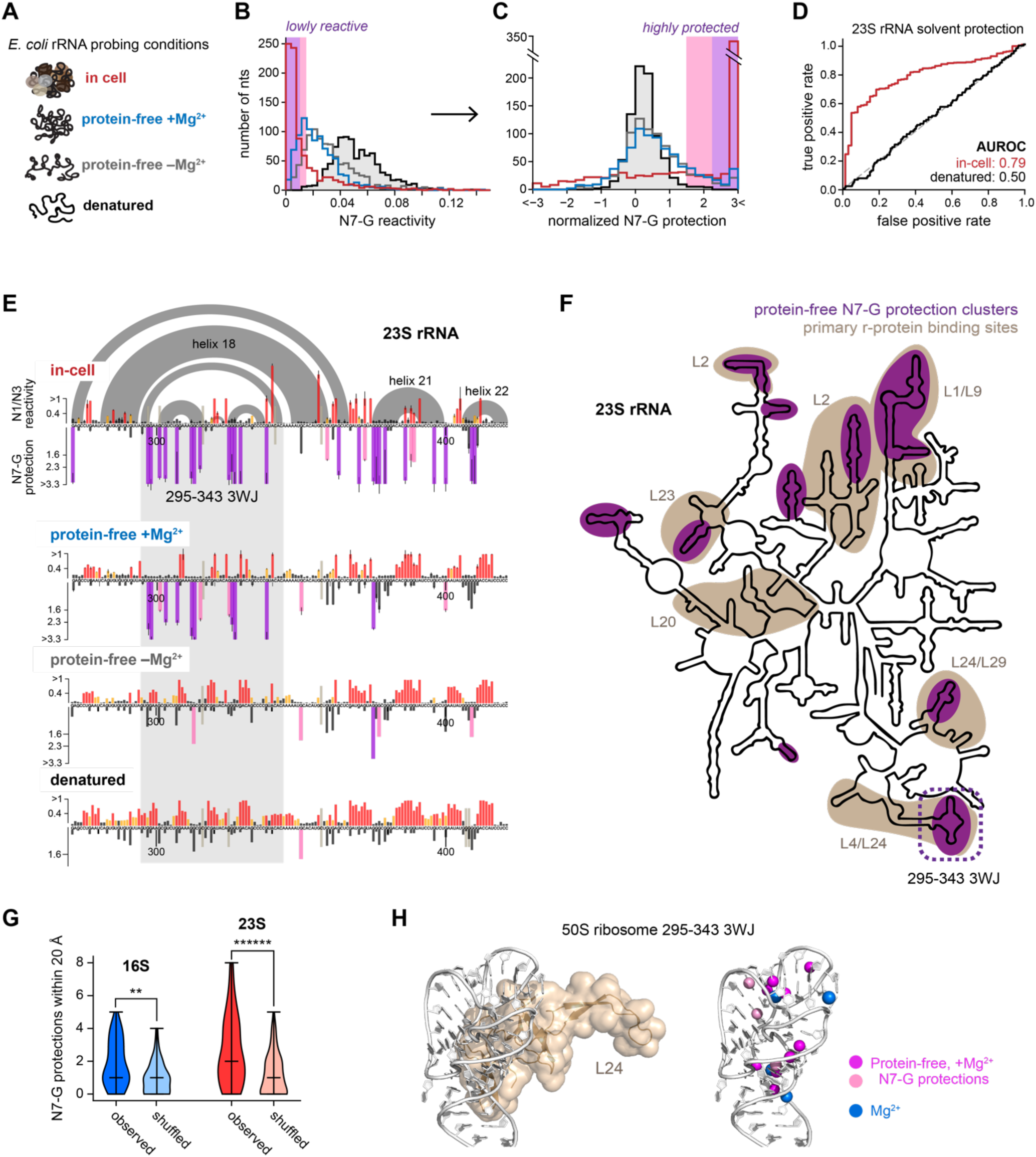
msDMS-MaP reveals self-folding tertiary modules in rRNA. (A) *E. coli* rRNA folding conditions probed by msDMS-MaP. (B, C) Distribution of N7-G (B) modification rates and (C) normalized protection scores at each folding condition. Pink and purple backgrounds denote significantly protected (lowly reactive) values. (D) Receiver operator characteristic (ROC) curves quantifying the ability of N7-G protections to discriminate between solvent inaccessible versus accessible positions in the assembled 50S ribosome. (E) Normalized N1/3 and N7 reactivity profiles for a segment of the 23S rRNA. In-cell and protein-free+Mg^2+^ values represent the average over two biological replicates, with error bars denoting the standard deviation. Protein-free-Mg^2+^ and denatured data are from single experiments. Gray arcs at top indicate base pairing interactions. The 295-343 3WJ is highlighted. (F) Map of self-folded tertiary domains in 23S rRNA identified from N7-G protections under protein-free+Mg^2+^ conditions (purple) and binding sites of primary r-proteins (brown). (G) Clustering of protected N7-G sites in three-dimensional space in protein-free rRNA versus if protections were randomly shuffled across G nucleotides. Distances between N7-G sites were computed using PDB 7K00^85^. Significance was computed using a two-sided Mann-Whitney U test; **, p<10^−3^, ******, p<10^−6^. (H) Left, structure of the 295-343 3WJ from 23S rRNA recognized by primary r-protein L24 (brown). Right, N7-G protections measured under protein-free+Mg^2+^ conditions and bound Mg^2+^ ions in purple and blue, respectively. Structures are excerpted from PDB 7K00^85^.

Inspired by the strategies used to analyze SHAPE^6,33^ and N1/N3 DMS^30,34^ experiments, we developed a normalization scheme to permit robust identification of protected N7-G sites across RNA systems and experimental conditions (Fig. 2C, E). Raw N7-G rates are adjusted for sequence context and log-transformed to obtain a protection score relative to the reactivity of an average unprotected guanine (Methods; Fig. S2G, H). Protection scores typically range between –3 and 3, with scores >0 indicating protection and scores <0 indicating enhanced reactivity (Fig. 2C). We selected protection scores of 1.6 and 2.3 as indicators of significant and very significant protection, respectively, corresponding to false discovery rates of 1% and 0% using the denatured rRNA as a null reference. We highlight protection scores meeting these thresholds in pink and purple, respectively.

Normalization of our in-cell rRNA data revealed that, as expected, most guanines are highly protected at the N7 position (Fig. 2C, E, S3, S4). These protected sites map out protein binding sites, tertiary interactions, and otherwise solvent inaccessible regions of the assembled ribosome. For example, the 295-343 three-way junction (3WJ) features strong protections in both base-paired and single-stranded regions deriving from RNA tertiary interactions and binding by proteins L4 and L24 (Fig. 2E). Neighboring helices 21 and 22 are bound by protein L28 and buried in the interior of the ribosome, respectively, and also feature strong N7-G protections. By comparison, the intervening helix 18 is solvent exposed and lacks protections (Fig. 2E, S2J). Overall, N7-G reactivity discriminates accessible versus inaccessible nucleotides with similar power as dedicated solvent accessibility probes (area under the receiver operating curve [AUROC] = 0.79 versus 0.75 for NAz ^12^; Fig. 2D).

Most rRNA N7-G protections disappear under protein-free conditions (Fig. 2C, E, S3, S4), consistent with most in-cell protections deriving from bound proteins and protein-dependent tertiary structures. However, some regions continue to exhibit N7-G protections, particularly in the presence of Mg^2+^ (Fig. 2C, E, F, S3, S4). Isolated regions of the *E. coli* 16S and 23S rRNAs have been reported to self-assemble into tertiary structures in the absence of proteins^35,36^. However, the difficulty of probing RNA tertiary structure has prevented systematic characterization of rRNA self-assembly. Protein-free N7-G protections cluster together on the rRNA secondary structure (Fig. S2I), and are also significantly clustered in three-dimensional space when mapped onto the structure of the assembled ribosome (Fig. 2G). This clustering is consistent with N7-G protections mapping out self-folding tertiary modules. Building on this observation, we used clustered N7-G protections to identify self-folding modules across the >4 kilobases of rRNA sequence (Methods). This strategy successfully identified known self-folding domains, such as 5’ subdomains of the 16S rRNA^37^, the 23S GTPase center^38^ and the L1 stalk^39^, as well as numerous previously unreported self-folding modules such as the 23S 295-343 3WJ (Fig. 2E, F, S3, S4). Visual inspection of the complete ribosome structure supports that these self-folding modules have self-contained tertiary structures (Fig. 2H, S5). Remarkably, these self-folding modules strongly coincide with binding sites of primary assembly proteins (Fig. 2F, H, S3, S4)^40,41^. Thus, our data are consistent with rRNA folding playing a central role in encoding ribosome assembly hierarchy^35,36^, with independently stable rRNA subdomains providing specific binding interfaces that recruit primary proteins to nucleate the assembly process. More broadly, these rRNA experiments establish the value of msDMS-MaP for identifying complex tertiary structures that are largely invisible to conventional probing methods.

### N7-G protections identify protein binding sites and tertiary structures across diverse systems

We next validated msDMS-MaP on a broader collection of RNA systems, including RNAs folded in solution and RNP complexes in living bacterial and human cells (Fig. 3A). N7-G protections demonstrate an average positive predictive value (PPV) of 91% and sensitivity (Sens) of 38% in identifying solvent protected guanines observed in high-resolution structures (Fig. 3A). Further analysis revealed that many “false positive” N7 protections are adjacent to unresolved protein chains, suggesting that they reflect true protections in solution. Conversely, many N7-reactive but solvent inaccessible sites (false negatives) are only “protected” by the RNA helical backbone and are likely exposed by helical flexing in solution. Thus, these PPV and Sens metrics likely underestimate the true specificity and sensitivity of N7-G protections for identifying guanines involved in complex structures.

**Figure 3:**
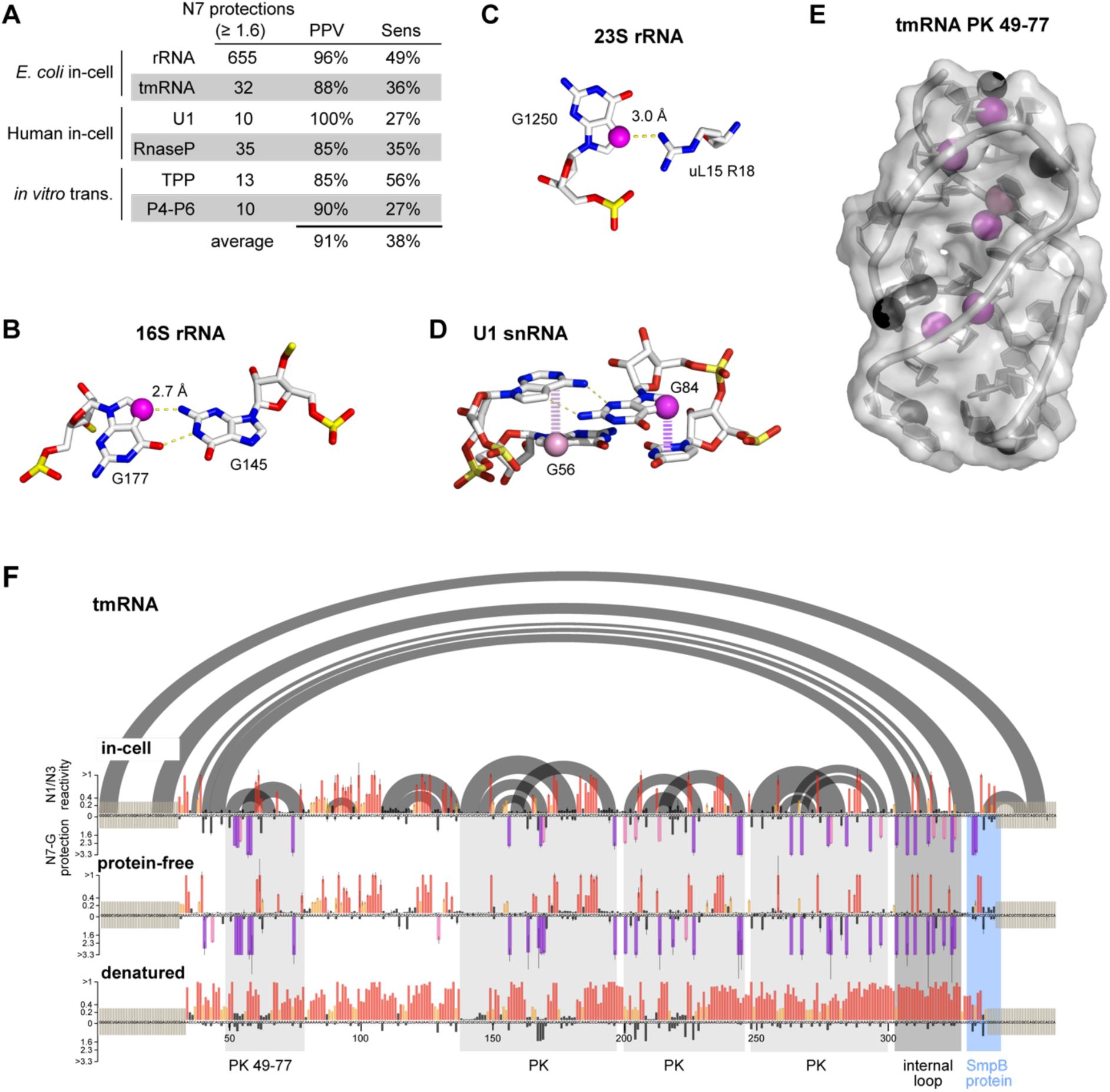
N7-G protections accurately report on diverse RNA tertiary and quaternary interactions. (A) Number of N7-G protections observed in different RNA and RNP systems, positive predictive value (PPV), and sensitivity (Sens) of these protections in identifying solvent protected nucleotides. Values represent averages over two replicates for each RNA. (B) Example of N7-G site (purple) protected by a Hoogsteen base-pairing interaction (G177 in E. coli 16S rRNA; PDB 7K00^85^). (C) Example of N7-G site protected by hydrogen bonding with a protein (G1250 in E. coli 23S rRNA; PDB 7K00^85^). (D) Example of N7-G protections from atypical base-stacking interactions (G84 and G56 in U1 snRNA; PDB 6QX9^93^). (E) Example of N7-G protections due to solvent inaccessibility in pseudoknot (PK) 49-77 from *E. coli* tmRNA (PDB 7ABZ^42^). Protected and unprotected N7-G positions are shown in purple and black, respectively. (F) Normalized msDMS-MaP data for *E. coli* tmRNA probed in cells and under protein-free and denatured conditions. Gray arcs at top indicate base pairing interactions. PKs and tertiary interactions are highlighted below in gray, and SmpB protein binding site in blue. For in cell and protein-free conditions, reactivities represent the averages of two replicates with error bars denoting the standard deviation. Denatured data is from a single experiment.

N7-G protections derive from three different types of molecular interactions. First, direct hydrogen bonding of N7-G with other RNA nucleotides in the context of non-canonical pairs or tertiary interactions, or with proteins or other bound ligands, conveys strong protection from DMS modification (Fig. 3B, C). Second, atypical stacking interactions that involve greater than normal overlap between the N7 and adjacent bases can provide moderate-to-strong N7-G protection even when the N7 position is otherwise solvent accessible (Fig. 3D). These stacking protections are observed in structured loops and non-canonical base-pairs and likely arise from altered nucleobase electronics or steric occlusion of the N7 site. Third, large tertiary structures and protein-RNA interfaces can bury N7-G sites, making them inaccessible to DMS even if they are not participating in direct interactions (Fig. 3E). Large tertiary structured domains and RNA-protein interfaces typically feature clusters of strong protections that arise from all three mechanisms. By comparison, isolated N7-G protections reflect local tertiary structures, such as ordered internal loops, or smaller RNA-protein interfaces.

The *E. coli* non-coding RNA tmRNA provides a vivid example of how N7-G protections illuminate complex RNA features (Fig. 3F). tmRNA has an intricate fold consisting of a three-way junction bound by the protein SmpB, a structured internal loop, and four pseudoknots^42^. These structures all feature low N1/N3 DMS reactivities, but distinguishing these complex structures from basic secondary structures is challenging. By comparison, each complex structure is clearly delineated by N7-G protections. N7-G sites in pseudoknots are solvent-protected by loops packing into the grooves of knotted helices (Fig. 3E, F). N7-G sites in structured internal loops and junctions are protected by RNA-RNA and RNA-protein hydrogen bonding, atypical stacking, and burial from solvent mechanisms. Under extracted, protein-free conditions, pseudoknots and structured loops feature even stronger N7-G protections, consistent with these structures folding more stably outside of cells (Fig. 3F)^43^. Conversely, N1/3 and N7-G reactivity increases at the SmpB binding site, supporting this as a protein binding site (Fig. 3F). These data further emphasize the ability of msDMS-MaP to reveal diverse, functionally important motifs that are challenging to identify by conventional probing.

### Correlated N7-G modifications report on coupled tertiary folding and protein binding events

msDMS-MaP experiments also permit measurement of correlated chemical modifications in single RNA molecules, which can provide powerful information about higher-order RNA structure and dynamics^44^. RNA interaction group (RING) analysis is a single-molecule strategy for identifying pairs of nucleotides that are significantly co-modified together (Fig. 4A)^7,44^. In standard N1/N3 DMS-MaP datasets, RINGs reveal secondary structure base pairs and three-dimensional structural communication such as tertiary interactions. We therefore explored whether N7-G modifications also generate RING signals that inform on RNA architecture. Indeed, RING analysis of our msDMS-MaP datasets revealed dense networks of RINGs between N1/3 and N7-G sites, and, less commonly, between pairs of N7-G sites (Fig. 4, S6).

**Figure 4:**
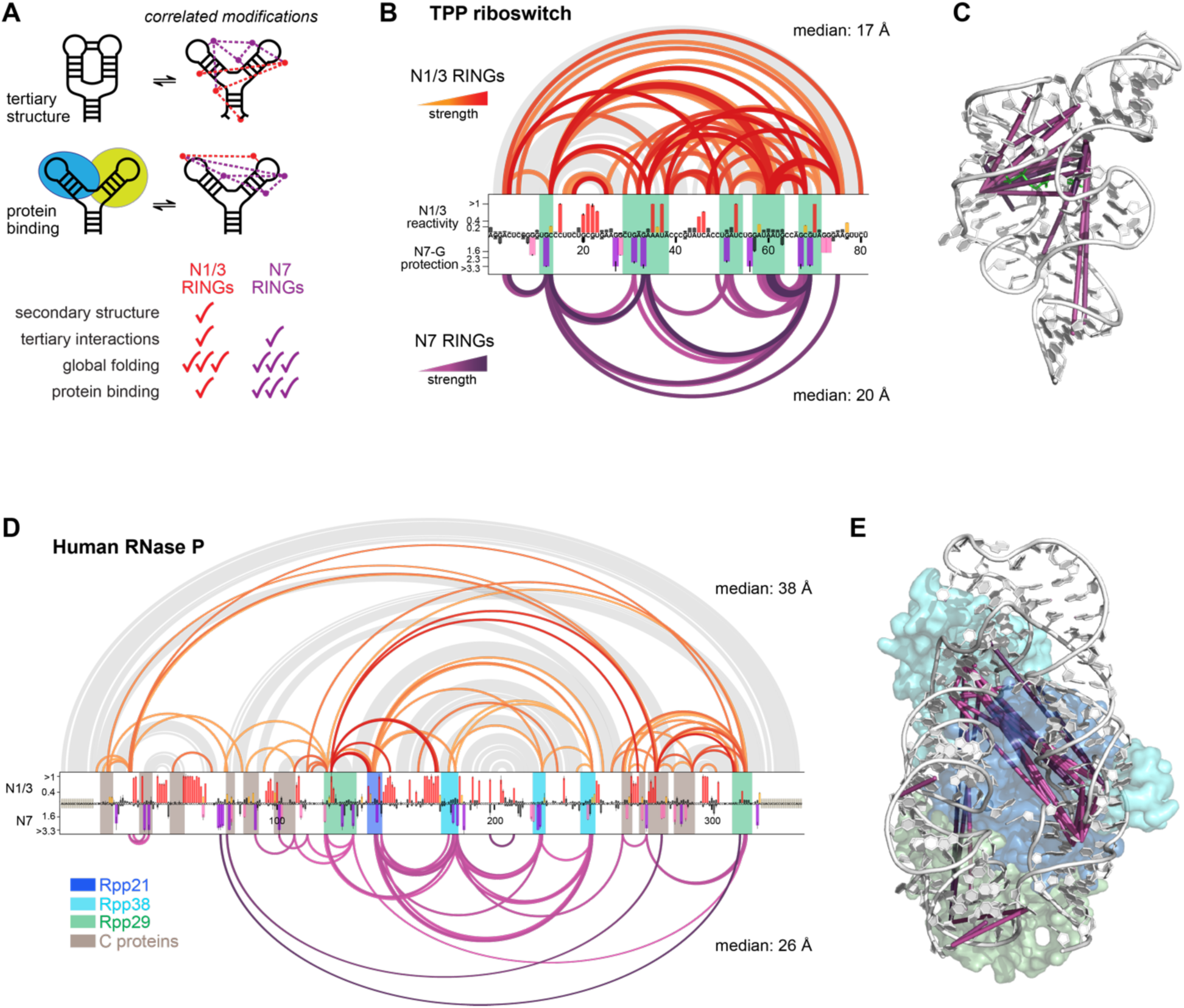
N7-G RINGs identify cooperative tertiary folding and protein binding domains. (A) Mechanism of correlated N1/3 and N7 modification, generating RNA interaction groups (RINGs). (B) Conventional N1/3-to-N1/3 RINGs (top, red) and N7-to-N1/3 and N7-to-N7 RINGs (bottom, purple) measured in ligand bound TPP riboswitch. Darker colors indicate stronger RINGs. RINGs were required to be observed in two independent replicates. N1/3 and N7-G reactivity profiles represent the average of two replicates, with error bars denoting the standard deviation. Gray arcs at top indicate base pair interactions. Nucleotides comprising the ligand binding pocket and tertiary interactions are highlighted in green. Median three-dimensional distances spanned by N1/3 and N7 RINGs are listed at top and bottom, respectively. (C) N7-to-N1/3 and N7-to-N7 RINGs from (B) projected onto TPP riboswitch three-dimensional structure (PDB 2GDI^32^). (D) RINGs measured in human RNase P probed in living RPE-1 cells. Darker colors indicate stronger RINGs. RINGs were required to be observed in two independent replicates. N1/3 and N7-G reactivity profiles represents the average of two replicates, with error bars denoting the standard deviation. Gray arcs at top indicate base pair interactions. Median three-dimensional distances spanned by N1/3 and N7 RINGs are listed at top and bottom, respectively. (F) N7-to-N1/3 and N7-to-N7 RINGs projected onto the specificity domain of human RNase P (PDB 6AHR^47^). RINGs and proteins are colored as in (D).

In RNAs with tertiary structures, N7-G RING networks report on coupled tertiary folding domains. For example, 4 N7-to-N7 and 22 N7-to-N1/3 RINGs are observed in the ligand-bound TPP riboswitch (Fig. 4B). These RINGs predominantly span across structured internal loops and across the TPP ligand binding site, reflecting N7-G sites and N1-A and N3-C sites that are coordinately protected by tertiary structure (Fig. 4B, C). RINGs are also observed between N7-G sites and more distal N1-A and N3-C sites, reporting on secondary structure base pairs that are coupled to tertiary structure formation. These N7 RINGs complement the conventional N1/3-to-N1/3 RINGs that are also observed in msDMS-MaP datasets (Fig. 4B). Whereas N1/3-to-N1/3 RINGs reflect both secondary and tertiary interactions, N7 RINGs predominantly reflect tertiary structure (Fig. 4B). This pattern of N7 RINGs mapping out tertiary folding domains with equal-to-greater specificity than N1/3-to-N1/3 RINGs was also observed in the P4-P6 domain of the *Tetrahymena* intron (Fig. S6A). Thus, N7 RINGs enhance standard RING analysis, providing rich information about tertiary folding that can be used for applications such as structure discovery and three-dimensional structure modeling^7^.

In RNAs that form quaternary complexes with proteins, N7-G RINGs also report on correlated protein binding events that occur at distal RNA sites. For example, RING analysis of in-cell probed human U1 snRNA revealed numerous N7-to-N1/3 and N7-to-N7 RINGs linking the binding sites of the 70K and Sm proteins, consistent with the known cooperative assembly of these proteins onto U1 (Fig. S6B, C)^45^. By comparison, few RINGs emanate from the U1A protein binding site, consistent with U1A binding independently^14,46^. RING analysis of human RNase P RNA also revealed a dense network of N7-to-N1/3 and N7-to-N7 RINGs spanning the binding interface of the Rpp21-Rpp29-Rpp38 protein heterotrimer (Fig. 4D, E). These RINGs report on burial of N7-G sites by the protein trimer with coupled folding of the RNase P specificity domain that brings correlated N7 and N1/3 sites into three-dimensional proximity (Fig. 4E)^47^. Binding sites for other RNase P proteins in the catalytic domain feature comparatively few RINGs, which may be explained by these proteins binding earlier and more stably than Rpp21-Rpp29-Rpp38^48,49^. Notably, the coupling between Rpp21-Rpp29-Rpp38 sites is challenging to discern by standard N1/3-N1/3 RING analysis, which reports a more complex mixture of direct and indirect secondary and tertiary correlations (Fig. 4D). Collectively, these data support that N7-G RINGs provide a sensitive and specific measure of coupled protein binding and tertiary folding events in living cells.

### msDMS-MaP resolves state-specific tertiary structures and protein binding sites in multi-state structural ensembles

Single-molecule DMS probing has also emerged as an impactful methodology for resolving multi-state RNA conformational ensembles in living cells^7,44,50–52^. RNAs that co-exist in multiple structures exhibit distinct patterns of co-modifications as nucleotides become coordinately reactivity versus unreactive. These correlated modification patterns can be deconvolved via maximum likelihood methods into multiple distinct underlying reactivity profiles (Fig. 5A). While powerful, existing deconvolution strategies typically can only resolve secondary structure ensembles. Many RNA and RNP conformational ensembles also feature coupled changes in tertiary and quaternary structure^3^, and measuring these coupled changes in structure remains difficult. We reasoned that single-molecule msDMS-MaP was uniquely suited for resolving such higher-order structural changes and therefore adapted the DANCE^52^ deconvolution framework to concurrently resolve N1/N3 and N7 reactivity profiles for multi-state RNA conformational ensembles (Fig. 5A).

**Figure 5:**
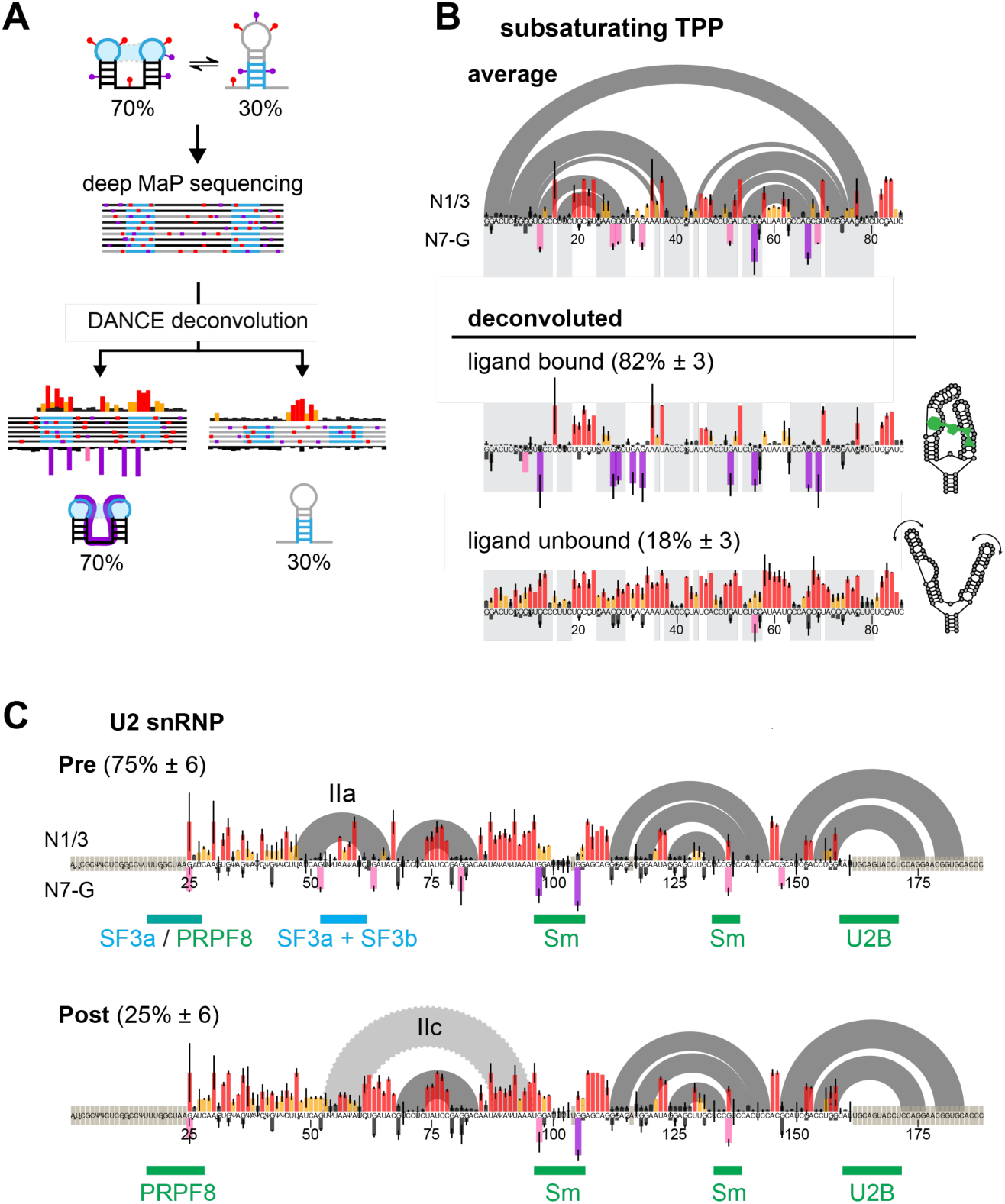
msDMS-MaP combined with DANCE resolves state-specific N7-G protections in RNA structural ensembles. (A) Schematic of DANCE strategy for deconvoluting N1/3 and N7-G reactivity profiles for multi-state RNA structural ensembles. (B) Averaged (top) and DANCE-deconvoluted (bottom) N1/3 and N7-G reactivity profiles for the TPP riboswitch probed under sub-saturating ligand concentrations (100 nM TPP ligand concentration). DANCE deconvolutes the ensemble into expected ligand-bound and unbound states. Reactivities and populations represent averages over three replicates, with errors bars and intervals denoting standard deviations. (C) DANCE-deconvoluted reactivity profiles of U2 snRNA in RPE-1 cells, corresponding to Pre-and Post-catalytic U2 snRNP states. Gray highlights indicate major structural changes in both states of U2 snRNA, and horizontal green and blue bars denote known protein binding sites. Population averages and standard deviations are calculated from three replicates.

To validate our N7-resolved DANCE strategy, we performed msDMS-MaP experiments on the P4-P6 and TPP riboswitch RNAs under subsaturating Mg^2+^ and TPP ligand concentrations, respectively, where these RNAs exist in dynamic equilibrium between tertiary folded and unfolded states^7,53,54^. Averaged analysis revealed increased N1/N3 reactivities and weakened N7-G protections compared to saturated folding conditions, supporting that the magnitude of N7-G protection provides a physically meaningful measure of average nucleotide accessibility (Fig. 5B, S7A). DANCE analysis successfully resolved both datasets into two-state ensembles consistent with the expected tertiary folded versus unfolded equilibrium (Fig. 5B, S7A). For both RNAs, N7-G protections specifically appear in the tertiary folded state and strengthen significantly compared to averaged values, matching protection strengths observed under saturated folding conditions (Fig. 5B versus 4B; Fig. S7A). Thus, msDMS-MaP combined with DANCE can accurately identify tertiary structured substates in complex structural ensembles.

We also evaluated the ability of msDMS-MaP to resolve RNA-protein quaternary ensembles using the human U2 snRNP as a model system. Genetics and high-resolution structures have shown that the U2 snRNA adopts multiple intra- and inter-molecular secondary structures that feature distinct protein interactions during the splicing cycle^55^. DANCE analysis of in-cell msDMS-MaP experiments resolved U2 into two conformational macrostates that primarily differ by the presence versus absence of stem IIa as assessed by N1/N3 reactivity (Fig. 5C). Stem IIa is a defining feature of pre-catalytic U2 complexes (free snRNP, and A to B^act^ spliceosome complexes)^55–57^, and thus we name the predominant IIa-containing state as **Pre**. The second minority state recapitulates the expected features of catalytic and post-catalytic complexes (B* to ILS spliceosome complexes), which feature a dynamic IIc stem in place of IIa (Fig. 5C)^55,58,59^. We thus name this minority state **Post**. Significantly, the **Pre** and **Post** states also exhibit distinct patterns of N7-G protections. The Pre state features multiple N7-G protections within the IIa stem, matching the known interactions with the SF3a and SF3b protein complexes that stabilize IIa (Fig. 5C)^55,56^. These protections are lost in the Post state, matching the expected detachment of the IIa region and heterogeneity amongst post-catalytic substates^55,58,59^. As a positive control, both Pre and Post states feature strong N7-G protections at the constitutive Sm protein binding site. The Pre/Post equilibrium and Sm-specific N-7G protections disappear when U2 is probed under protein-free conditions (Fig. S7C), confirming that protections observed in cells report on bound proteins. In sum, msDMS-MaP permits direct measurement of coupled RNA folding and protein binding events in RNA and RNP ensembles.

### msDMS-MaP reveals quaternary architecture of 7SK RNP ensemble

We sought to use msDMS-MaP to obtain an improved model of the conformational ensemble populated by the human 7SK snRNP in cells. 7SK is a highly conserved 331-334 nucleotide long RNA that regulates the activity of positive transcription elongation factor b (P-TEFb; also known as CDK9-Cyclin T)^60,61^. 7SK exists in multiple states that either bind and inhibit P-TEFb, or release P-TEFb to permit phosphorylation of RNA Pol II and licensing of transcription elongation. We previously showed using DMS-MaP and DANCE analysis that 7SK populates three distinct structural states in cells, which we termed A, B, and H, and which we assigned to P-TEFb bound, released, and putative intermediate states, respectively^52^. However, our experiments were unable to resolve any protein binding sites on the 7SK RNA, even though numerous proteins are known to interact with and presumably drive exchange between different 7SK states^61–67^. More broadly, the global quaternary architecture of 7SK snRNPs remains unknown^61^.

We acquired three in-cell replicates of single-molecule msDMS-MaP data from human RPE1 cells and two additional replicates from HeLa cells (Fig. 6A; Fig. S8A). DANCE analysis resolved N1/N3 data into three states that closely corresponded to the previously identified A, B, and H structures^52^. However, in a modest revision to our prior model of state A, our improved N1/N3 data support that state A features an extended SL1 stem in place of the SL0 stem observed in state B, consistent with cryo-EM structures (Fig. 6D; S8B, C)^68^.

**Figure 6:**
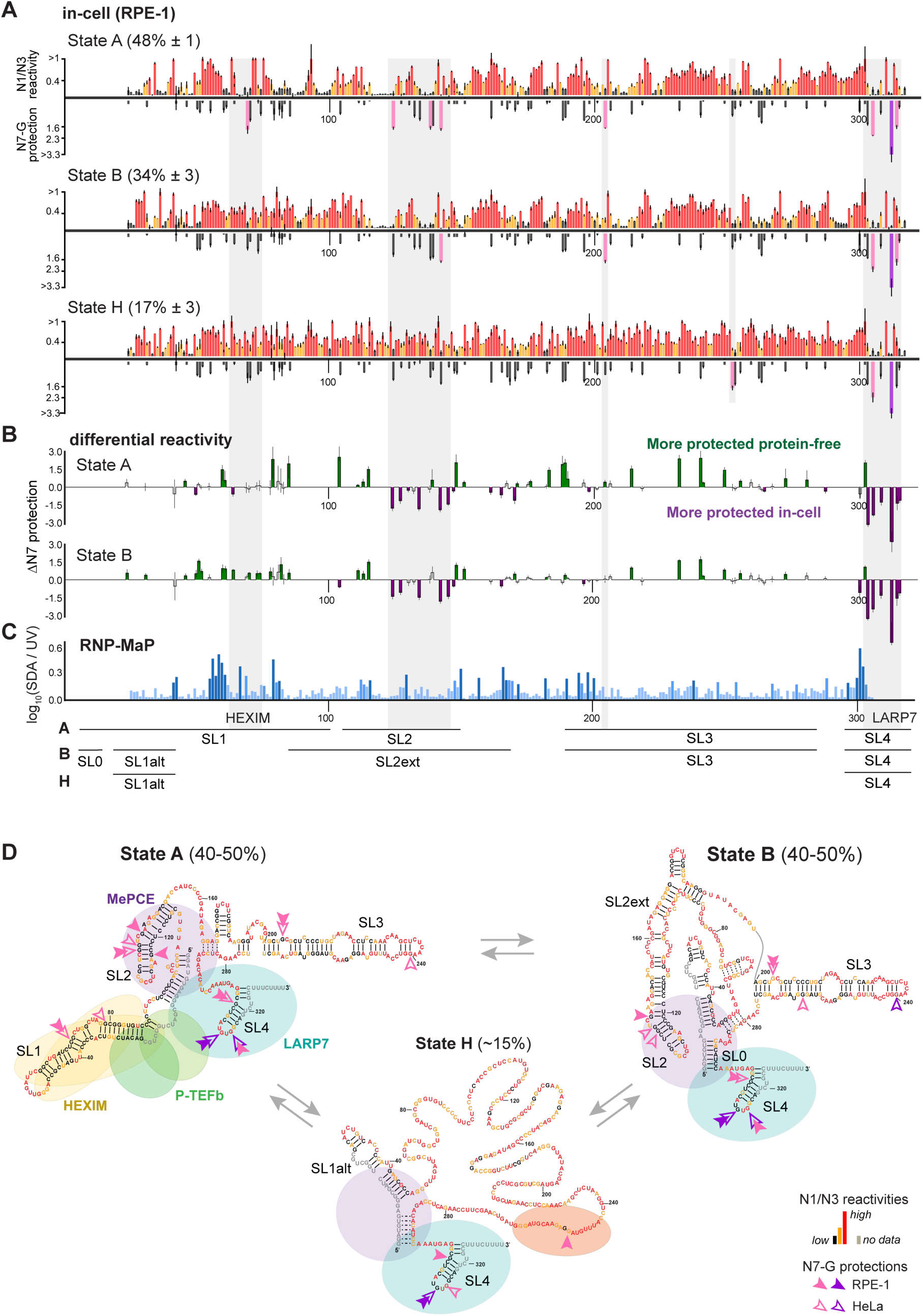
msDMS-MaP reveals quaternary architecture of the 7SK snRNP structural ensemble. (A) N1/3 and N7-G reactivity profiles for three-state 7SK snRNA structural ensemble resolved by DANCE, probed in human RPE-1 cells. Population averages and standard deviations are calculated from three replicates. (B) Differential analysis of N7-G protection in cells versus protein-free 7SK, showing State A (top) and State B (bottom). Differences represent averages between three in-cell and three protein-free replicates, with error bars denoting standard deviations. Nucleotides exhibiting statistically significant (p<0.05) increases in N7-G protection in cells are shown in purple, and statistically significant increases in protein-free conditions in green. (C) RNP-MaP analysis of 7SK RNA performed in HEK293T cells. Data represent the average of two replicates. RNP-MaP sites exceeding empirical significance thresholds^14^ are highlighted in green. (D) Revised model of the 7SK structural ensemble in cells. Consensus models developed in Ref. 52 were manually adjusted based on msDMS-MaP data (e.g. removal of SL0 in State A). Possible unstable pairs are indicated with dashes. Nucleotides are colored by average in-cell msDMS-MaP N1/3 reactivity. N7-G protections observed in cells are shown with pink and purple arrows.

N7 reactivity analysis revealed distinctive patterns of N7 protections in each 7SK state, consistent with global remodeling of quaternary structure (Fig. 6A, S8A). As a positive control, strong protections are observed in all three states at SL4, the binding site for the constitutive protein LARP7^68,69^. By comparison, state A features unique protection at G69, precisely corresponding to the expected binding site of the protein HEXIM1 that bridges P-TEFb to 7SK^70–72^. Interestingly, states A and B both feature N7 protections in SL2, a highly conserved domain of unknown function^61^. We also observe several N7-G protections in the SL3 region, including at G205 in states A and B and at G252 in the unstructured H state (Fig. 6A). The G252 protection provides evidence that SL3 unfolding in state H is stabilized by bound proteins. However, the G252 protection is not observed in HeLa cells (Fig. S8A), suggesting that protein occupancy at this site varies based on cell type and state.

To assess whether these in-cell N7 protections reflect bound proteins, we collected msDMS-MaP datasets on extracted, protein-free 7SK RNAs (Fig. S9). Protein-free data closely matched previously published measurements and clustered into three states, corresponding to A, B, and a state termed M that is exclusively observed under protein-free conditions (Fig. S9B)^52^. N7-G protections consistent with isolated non-canonical base-pairs are observed in each of these states. Notably, SL1 of state A features multiple N7-G protections, consistent with NMR and X-ray crystallography studies showing that SL1 forms extensive local tertiary interactions *in vitro* (Fig. S9A)^70,71,73^. Differential analysis of in-cell versus protein-free state A revealed several small but statistically significant increases in N7-G protection in SL1, exactly overlapping the expected binding site of HEXIM/P-TEFb (Fig. 6B). Differential analysis also revealed significant in-cell-specific protections at SL2 and SL4 in states A and B, providing strong evidence that these are indeed protein binding sites (Fig. 6B). These in-cell protections are difficult to discern by N1/N3 differential analysis, which instead predominantly reports on destabilization of 7SK secondary structure in cells (Fig. S9B). In contrast to SL1, SL2, and SL4, N7 protection at G205 in states A and B is increased in protein-free RNA, indicating that the G205 protections observed in cells reflects non-canonical pairing. Note that state H cannot be analyzed for differential protection because state H only exists inside of cells. As orthogonal validation, we also performed RNP-MaP experiments^14^, which use RNA-protein cross-linking to directly map protein binding sites on RNA inside cells; these RNP-MaP data support protein binding at regions identified by msDMS-MaP (Fig. 6C).

We integrated these data to develop a refined model of 7SK quaternary structure (Fig. 6D). Our data provide strong evidence that P-TEFb exclusively binds state A in cells. However, this binding appears dynamic, with much of the expected HEXIM-P-TEFb binding interface demonstrating only moderate N7 protection. N7 protections observed in SL2 across states A and B are most consistent with interaction of this region with core snRNP proteins MEPCE or LARP7. Notably, *in vitro* reconstitution experiments also reported evidence of interactions between SL2 and MEPCE or LARP7^67,74^. Finally, protections in the SL3 region in the unstructured state H support that this state is a product of protein-induced unfolding. Multiple P-TEFb “release factors” have been implicated as binding this SL3 region, and most of these are RNA helicases and single-stranded binding proteins^61,62,65,66,75–77^. These data are thus suggestive that state H represents an intermediate along the P-TEFb release pathway. Overall, our data emphasize the close relationship between 7SK RNA structural and protein compositional remodeling and highlight the power of msDMS-MaP for visualizing complex RNP dynamics.

## DISCUSSION

We introduced msDMS-MaP as a new strategy for simultaneously probing RNA secondary and higher-order structure. The core innovation of msDMS-MaP is using tautomerism to decode otherwise invisible N7-methylguanine (N7-G) modifications during reverse-transcription reactions. msDMS-MaP is thus substantially easier to perform than existing depurination methods for measuring N7-G modifications, yielding N7-G reactivity information essentially “for free” from a standard chemical probing workflow. msDMS-MaP analysis is automated and implemented in the widely used ShapeMapper analysis software(Fig. S10)^78^, further facilitating easy adoption. The scalable msDMS-MaP workflow enabled us to perform the first comprehensive investigation of DMS N7-G reactivity, including in living bacterial and human cells. Combined, our data establish N7-G protection as a sensitive and specific measure of solvent inaccessibility, revealing otherwise imperceptible RNA tertiary structures and protein binding sites that underpin RNA function.

msDMS-MaP offers several advantages over alternative methods for probing higher-order RNA structure. msDMS-MaP experiments are comparatively easy to perform, can be performed in cells, require only inexpensive and widely available reagents, and are compatible with most high-throughput sequencing library preparation protocols. msDMS-MaP is also uniquely capable of measuring coupled changes in secondary, tertiary, and quaternary structure across multi-state RNA ensembles. However, msDMS-MaP can only probe higher-order structure of G nucleobases. Because N7-G modifications are decoded with only ∼20% efficiency, msDMS-MaP experiments further require high overall levels of DMS modification for reliable quantification, which may perturb some RNA molecules.

A key application enabled by msDMS-MaP is improved discovery of RNA tertiary structures. Tertiary structures are particularly likely to have significant biological functions. They are also more likely to contain specific small molecule binding pockets, making them high-value targets for RNA-targeted drug discovery^79,80^. We show that diverse tertiary structures, ranging from internal loops to pseudoknots, generate strong N7-G protections that permit confident identification of these motifs from commonplace secondary structures. In many cases, these structures also feature dense networks of correlated N7-G and N1/N3 modifications (RINGs) that define cooperative folding domains. Our studies of protein-free *E. coli* ribosomes highlight the potential of msDMS-MaP as a discovery tool, enabling us to identify self-folding tertiary domains across >4,000 nucleotides of RNA sequence. These self-folding domains, most of which were previously uncharacterized, coincide with binding sites of primary ribosome assembly proteins, supporting that ribosome assembly hierarchy is encoded by rRNA tertiary folding potential.

Our studies of endogenous human U1, U2, and RNase P snRNPs also demonstrate msDMS-MaP as a powerful strategy for resolving RNA-protein interactions. N7-G protection scales with protein binding occupancy, with strong protections signifying high occupancy whereas weaker protections indicate weaker or more dynamic protein binding. Correlated N7-G RING signals further reveal cooperative protein binding events that span across distal RNA sites, while ensemble deconvolution can be used to determine protein binding to distinct RNA structural states. At present, distinguishing whether N7-G signals reflect bound proteins versus tertiary structure requires comparisons to matching protein-free datasets. The identity of bound proteins also must be determined via orthogonal methods. Nonetheless, the ability of msDMS-MaP to immediately reveal putative quaternary interactions will accelerate discovery and characterization of functional RNA elements in living cells.

The 7SK snRNP exemplifies the challenges of resolving RNP mechanisms, with prior analyses indicating that 7SK exists in multiple structures and varied protein compositions, but with the relationship between these structural and compositional dynamics being unclear^52,61,67,81^. By providing an integrated view of the 7SK ensemble in cells, our msDMS-MaP study establishes that remodeling of 7SK RNA secondary structure is closely coupled to quaternary structure remodeling. This coupled remodeling is reminiscent of spliceosome snRNP mechanisms^82^ and likely reflects a more general mode of how non-coding RNAs function. Our data also provide new insights into 7SK global architecture, including likely interactions of SL2 with core proteins MEPCE and LARP7, and that 7SK unfolding in the heterogenous H state is associated with protein binding to SL3. Better defining the protein composition and function of H state represents a key goal for future studies.

In sum, msDMS-MaP is a powerful tool for discovering, prioritizing, and characterizing functional RNA motifs. While not pursued here, we further anticipate that N7-G reactivity information will facilitate improved RNA secondary and three-dimensional structure modeling^7,15,83^. The ability to measure both N1-G and N7-G reactivities suggests that msDMS-MaP will also be particularly well-suited for defining G-quadruplex structures^22,84^. Finally, we suggest that tautomeric stabilization may represent a more general strategy for resolving other types of RNA chemical modifications, both in the context of chemical probing experiments and for profiling endogenous RNA modifications.

## ACKNOWLEDGEMENTS

We thank J. Meehan (BCM) for feedback on the method and manuscript. We thank Kevin Weeks and Patrick Irving (UNC) for helpful discussions and ongoing support of the ShapeMapper code base. We thank Anna Pyle and Li-Tao Guo (Yale) and the Recombinant Protein Production and Characterization Core at BCM for help with MarathonRT production. This work was funded by the National Institutes of Health [R35 GM147010 to A.M.M.] and the Cancer Prevention and Research Institute of Texas [RR190054 to A.M.M.]. A.M.M. is a CPRIT Scholar in Cancer Research and a Beckman Young Investigator.

## CONFLICT OF INTEREST STATEMENT

A.M.M. is an advisor to and holds equity in RNAConnect, Inc., which holds patent rights to MarathonRT. The research in this paper was conceived and funded independently of RNAConnect. A.M.M. has also consulted for Ribometrix.

## METHODS

### Optimization of Marathon RT conditions

MarathonRT optimization experiments were performed using total *E. coli* RNA extracted and treated with ethanol following the protocol described below for protein-free *E. coli* probing experiments. RT primers were designed targeting ∼150 bp regions encompassing the 16S 527 and 23S 2069 natural m7G bases (Table S1)^85^. MarathonRT reverse-transcription reactions were adapted from published protocols^86^. 1 μg of total RNA was mixed with 1 μL 10 mM dNTPs and 1 μL 5 μM RT primer, adjusted to 6.8 μL with H_2_O, denatured at 65 °C for 10 min, and then cooled at 4°C for 2 min. 12.2 μL of 1.64× reaction buffer (variable) was then added, followed by incubation at 23 °C for 2 min, addition of 1 μL of Marathon enzyme (20 U; Kerafast), incubation at 42°C for 3 hr, and heat inactivation at 95°C for 1 min. For pH variation experiments, 1× buffer consisted of 50mM Tris-HCl (pH 7.5 – 9), 200 mM KCl, 5 mM DTT, 1 mM MnCl_2_, and 20% glycerol. For Mn^2+^ variation experiments, 1× buffer consisted of 50mM Tris-HCl (pH 8.3), 200 mM KCl, 5mMM DTT, 0.5 – 3 mM MnCl_2_, and 20% glycerol. Following reverse-transcription, cDNA was purified using SPRI beads (AMPure XP [Beckman Coulter]). Sequencing libraries were prepared following the two-step PCR strategy^87^.

### DMS probing of *in vitro* transcripts

DNA templates were purchased as gBlocks (IDT; Table S1), PCR amplified (Q5 DNA polymerase; NEB) and purified (PureLink PCR column; Invitrogen). Run-off transcription was then performed using 5 ng of PCR amplified template (HiScribe T7 High Yield RNA Synthesis kit; NEB), followed by DNase treatment (TURBO DNase; Invitrogen) and purification by SPRI beads (Mag-Bind TotalPure NGS [Omega Biotek] or AMPure XP [Beckman Coulter]). RNA size and purity were confirmed via TapeStation analysis (Agilent).

For TPP experiments shown in Figures 1 and 5, 30 pmol RNA in 30 µL H_2_O was denatured at 95°C for 2 min followed by snap-cooling on ice for 2 min. The RNA was refolded at 37°C by adding 18 µL of 3.2 folding buffer (1x folding buffer: 200 mM bicine (pH 8.0), 200 mM potassium acetate (pH 8.0), and 10 mM MgCl_2_) and incubating at 37 °C for 15 min. 6 µL of TPP ligand (variable concentration) or H_2_O was then added and the RNA incubated for an additional 15 min at 37 °C. 9 µL of each reaction were transferred to a separate tube and treated with 1 µL of 1.7 M DMS solution in ethanol or 1 µL of ethanol for 6 min at 37 °C. The reaction was quenched with an equal volume of 20% 2-mercaptoethanol, and the RNA was purified by ethanol precipitation.

For TPP experiments shown in Figure S1 and 4, 4 µg of TPP RNA in 20 µL H_2_O was refolded at 37 °C by adding 25 µL of TPP ligand folding buffer (1X buffer: 200 mM bicine (pH 8.3), 1 µM TPP ligand, 200 mM potassium acetate (pH 8.0), 10 mM MgCl_2_) and incubating at 37 °C for 40 min. 20 µL of each reaction was transferred to a separate tube and treated with 4 µL of 1.5 M DMS in ethanol or 4 µL of ethanol for 6 min at 37 °C. The reaction was quenched with 100 µL of 20% 2-mercaptoethanol, and the RNA was purified (RNA Clean & Concentrator, Zymo). This second protocol uses a ∼50% higher DMS concentration and thus generates ∼40% higher modification rates. Normalized N7-G protection scores were highly reproducible across the two TPP probing protocols (R=0.92). The 1.5x DMS protocol produces a larger quantity of measurable RINGs due to the higher modification rates of these samples, but both protocols yield qualitatively similar N1/N3 and N7 RING patterns.

For P4-P6 experiments, 20 pmol RNA in 15 µl H_2_O was denatured at 95 °C for 2 minutes and snap-cooled on ice for 2 minutes. The RNA was then refolded at 23 °C by adding 12 µl of 2.25 folding buffer (1X folding buffer: 200 mM bicine (pH 8.0), 100 mM NaCl, varied MgCl_2_) and incubating for 30 min. 9 µL of folded RNA was transferred to a separate tube and treated with 1 µL of 1.7 M DMS solution in ethanol or 1 µL of ethanol for 6 min at 37 °C. The reaction was quenched with an equal volume of 20% 2-mercaptoethanol, and the RNA was purified by ethanol precipitation.

### DMS probing of living *E. coli* cells

DMS probing of living *E. coli* cultures was performed as described previously^18^. Briefly, an overnight culture of *E. coli* K-12 MG1655 was inoculated in LB and grown at 37 °C to OD_600_ ≈0.5. Cultures were pelleted at 4000 g for 5 min and resuspended in 1× bicine folding buffer [200 mM bicine (pH 8.3 at room temperature), 200 mM potassium acetate, 5 mM MgCl_2_]. Cells were then treated with 0.7 M DMS in ethanol for 6 min at 37°C or with ethanol for 6 min at 37 °C and the reaction was quenched with an equal volume of ice-cold 20% 2-mercaptoethanol. Cells were pelleted at 10,000 *g* for 5 min at 4 °C, treated with 1 mg/mL lysozyme for 5 min on ice, and then RNA isolated using TRIzol reagent (ThermoFisher).

### DMS probing of protein-free *E. coli* RNA

DMS probing of protein-free *E. coli* K-12 MG1655 total RNA was performed as previously described^18^. Briefly, overnight *E. coli* cultures were inoculated in LB and grown at 37 °C until OD_600_ ≈0.5, followed by cell pelleting and resuspension in lysis buffer [15 mM Tris– HCl (pH 8), 450 mM sucrose, 8 mM EDTA (pH 8), 0.04 mg/mL lysozyme]. RNA was extracted using 3x phenol/chloroform/isoamyl alcohol (PCA) followed by 3x chloroform and exchanged into 1× folding buffer [200 mM bicine (pH 8.3), 200 mM potassium acetate (pH 8.0), 0 or 5 mM MgCl_2_]. After equilibration at 37 °C for 10 min, 9 volumes of total RNA were treated with 1 volume of 1.7 M DMS solution in ethanol or 1 volume of 100% ethanol and incubated at 37°C for 6 min. The reaction was quenched with ten volumes of ice-cold 20% 2-mercaptoethanol, RNA purified (RNeasy Midi, Qiagen), DNase treated (TURBO DNase, ThermoFisher), and purified again (RNeasy Midi, Qiagen).

### DMS probing on denatured *E. coli* RNA

Overnight *E. coli* cultures were inoculated in LB and grown at 37 °C to OD_600_ ≈0.5. Cells were pelleted at 5000 *g* for 5 min at 23 °C, then resuspended in 1 mL of 1 mg/mL lysozyme in 0.5x TE (pH 8.0) and incubated on ice for 5 min. RNA was then extracted using 8 mL of TRIzol (Sigma-Aldrich) and quantified by UV absorbance (NanoDrop). 13 μg of RNA in 7 M urea was denatured at 98 °C for 1 min, cooled at 4 °C for 1 min. Samples were split evenly into two aliquots and treated with 1 volume 1.7 M DMS in ethanol or 1 volume of pure ethanol in 9 volumes of total RNA at 37 °C for 6 min. Samples were then quenched with 10 volumes of 2-mercaptoethanol on ice and purified by ethanol precipitation.

### DMS probing of RPE-1 and HeLa cells

hTERT-RPE1 cells were maintained in DMEM-F12 with HEPES (Gibco) with 10% FBS (Gibco), 100 U/mL Pen/Strep (Gibco), 2 mM sodium pyruvate (Gibco), and MEM non-essential amino acids (Gibco) at 37 °C and 5% CO_2_. HeLa cells were obtained from the Baylor College of Medicine Tissue Culture Core and maintained in DMEM with 10% FBS, 100 U/mL Pen/Strep (Gibco), 2 mM sodium pyruvate (Gibco), and MEM non-essential amino acids (Gibco) at 37 °C and 5% CO_2_. Cells were regularly tested for and confirmed to be free of mycoplasma.

In-cell probing experiments were performed as previously described^52^. 1.5 x 10^6^ cells were seeded in a 10 cm culture dish and grown to 70% confluency. Immediately prior to probing, the medium was replaced with 5.4 mL fresh media supplemented with 200 mM bicine (pH 8.3) and cells incubated at 37 °C for 3 minutes. Cells were then treated with 600 μL of 1.7 M DMS in ethanol or 100% ethanol for 6 minutes at 37 °C, followed by quenching with 6 mL of ice-cold 20% 2-mercaptoethanol. Cells were scraped, pelleted, and RNA purified using TRIzol (Ambion) or RNeasy Mini Kit (Qiagen). RNA was quantified by UV absorbance (Nanodrop) and integrity confirmed by TapeStation analysis.

### DMS probing of protein-free human RNAs

Cell-free probing experiments were performed as previously described^52^. RNA was extracted from 70% confluent RPE-1 cells using TRIzol reagent (Ambion) or RNeasy Mini Kit (Qiagen), Dnase treated (Turbo DNase; Invitrogen), purified with SPRI beads (Mag-Bind TotalPure NGS [Omega Biotek] or AMPure XP [Beckman Coulter]), and quantified by UV absorbance. 4 μg of RNA in 50 μL water was denatured at 98 °C for 1 min, snap-cooled at 4 °C for 1 min, and folded by addition of 50 μL of 2x folding buffer (1x folding buffer: 200 mM bicine (pH 8.0), 200 mM potassium acetate (pH 8.0), 5 mM MgCl_2_) and incubation at 37 °C for 30 min. Samples were split into two 45 μL aliquots and treated with either 5 μL of 1.7 M DMS in ethanol or 5 μL of 100% ethanol at 37 °C for 6 min. After treatment, samples were quenched with an equal volume of 20% 2-mercaptoethanol, placed on ice, and purified by ethanol precipitation.

### msDMS-MaP reverse transcription

For a 20 μL reaction, 1-3 μg of total RNA or 1-5 pmol of IVT RNA was mixed with 1 μL of 10 mM dNTPs, 1 μL of either 5 μM gene-specific primers or 5-25 ng/μL random nonamer primers, and adjusted to 6.8 μL with H_2_O. Primers were then annealed by incubating the mixture at 65°C for 10 min followed by 4°C for 2 min. 12.2 μL of 1.64x RT buffer (1x buffer: 50 mM Tris-HCl (pH 9), 200 mM KCl, 5 mM DTT, 2mM MnCl_2_, and 20% glycerol) was then added to the RNA-primer mix followed by a 2 min incubation at 23 °C and then addition of 1 μL of MarathonRT enzyme (20 U). RT reactions were incubated at 42°C for 3 hrs, then heat inactivated at 95°C for 1 minute. Marathon enzyme was either purchased from Kerafast or expressed and purified as described^86^ from the pET-6xHis-SUMO-MarathonRT (Addgene plasmid #109029; gift of Anna Pyle). Following RT, cDNA was purified using SPRI beads (Omega Bio-tek; 1.8x bead ratio).

For 7SK msDMS-MaP experiments, we used a shortened locked nucleic acid (LNA) primer to permit measurement of LARP7 binding to SL4 located near to the 7SK 3’ terminus (Table S1). This primer also incorporates a unique molecular identifier (UMI) to permit detection and removal of PCR duplicates. To maximize the RT reaction yield (and number of measured UMIs), we first performed rRNA depletion (NEBNext rRNA depletion, NEB) on 3 µg of total cellular RNA and then input the product into a RT reaction.

### Sequencing library construction

msDMS-MaP sequencing libraries for TPP, P4-P6, tmRNA, U1, U2, RNase P, and 7SK were prepared following the two-step PCR strategy ^87^. For TPP and P4-P6, one-fifth of the RT product was input into PCR1 and performed according to the program: 98 °C for 30 s, 10 cycles of (98 °C for 10 s, 64 °C for 20 s, 72 °C for 20 s), and 72 °C for 2 min. For RNaseP and tmRNA, one-fifth of the purified reverse transcription (RT) product was input into PCR1 following the program: 98 °C for 30 s, 15 cycles of (98 °C for 10 s, 68 °C for 20 s, 72 °C for 20 s), and 72 °C for 2 min. For U1, one-fifth of the purified RT product was input into PCR1 and performed according to the program: 98 °C for 30 s, 15 cycles of (98 °C for 10 s, 61 °C for 20 s, 72 °C for 20 s), and 72 °C for 2 min. For U2, one-fifth of the purified RT product was input into PCR1 and performed according to the program: 98 °C for 30 s, 15 cycles of (98 °C for 10 s, 65 °C for 20 s, 72 °C for 20 s), and 72 °C for 2 min. For 7SK, the complete volume of RT product was input into PCR1 following the program: 98 °C for 30 s, 10 cycles of (98 °C for 10 s, 66 °C for 20 s, 72 °C for 20 s), and a final extension at 72 °C for 2 min. PCR1 products of all RNAs were purified by SPRI beads (Mag-Bind TotalPure NGS beads; 0.8x ratio). 1-2 ng of the purified PCR1 product was then input into PCR2 following the program: 98 °C for 30 s, 10-14 cycles of (98 °C for 10 s, 65 °C for 30 s, 72 °C for 20 s), and 72 °C for 2 min. PCR2 products were purified by SPRI beads (Mag-Bind TotalPure NGS beads; 0.8x ratio), assessed by Tapestation analysis (Agilent) and sequenced on an Illumina MiSeq instrument using v2 (2×150 bp or 2×250 bp) or v3 (2×300 bp) chemistry.

rRNA libraries were prepared from randomly primed total RNA following the Nextera strategy^87^. Following reverse transcription, cDNA was converted into double-stranded DNA (dsDNA) using the NEBNext second-strand synthesis module (NEB) using a two hour incubation at 16 °C. dsDNA was purified by SPRI beads (Mag-Bind TotalPure NGS beads; 0.65x ratio). dsDNA was then tagmented with Nextera XT (Illumina) following the manufacturer’s protocol and purified by SPRI beads (Mag-Bind TotalPure NGS beads; 0.56x ratio). Libraries were sequenced on an Illumina MiSeq using 2 x 300 paired-end sequencing.

### msDMS-MaP data processing by ShapeMapper 2.3

We automated msDMS-MaP analysis in the widely used ShapeMapper software package. This updated version of ShapeMapper (v2.3) is available for download at https://github.com/Weeks-UNC/shapemapper2. ShapeMapper computes N7 protections and N1/3 reactivities concurrently, generating data visualizations for immediate analysis and formatted text files for downstream analysis, such as arcPlot visualization, RING, and DANCE analysis. msDMS-MaP processing is invoked by using the --dms and --N7 flags (Fig. S10). ShapeMapper 2.3 also includes several minor updates to improve data quality control.

N1/3 reactivities are calculated exactly as previously described^18^. Briefly, N1-A, N3-C, and N3-U modification rates are calculated as the rate of mismatch mutations at A, C, and U positions observed in DMS-modified samples, minus the rate of mismatch mutations observed in unmodified samples. N1-G modification rates are calculated considering only G®C and G®U mismatches. Modification rates are then normalized on a nucleotide-specific basis to obtain normalized reactivities that follow a common scale between 0 and ∼1.

N7-G modification rates (*r*_*N7,i*_) are calculated as the rate of G→A mismatch mutations observed in DMS-modified samples minus the rate of G→A mismatches observed in unmodified samples:

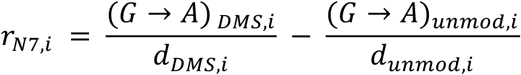

where (*G* → *A*)*_DMS,i_* and (*G* → *A*)*_unmod,i_* are the number of G→A mismatches and *d_DMS,i_* and *d_unmod,i_* are the sequencing depths of the DMS-modified and unmodified samples at position *i*.

Analysis revealed that N7-G modification rates depend systematically on sequence context, with guanines in 5’-GR contexts demonstrating ∼65% lower N7-G modification rates compared to 5’-GY contexts (R=purine, Y=pyrimidine; Fig. S2G). This sequence context bias was observed in all datasets, including RNAs probed under denaturing conditions. We thus conclude that this bias reflects differences in MarathonRT decoding efficiency, likely due to stacking with +1 purines stabilizing the non-mutagenic keto tautomer of m7G. This sequence dependence is removed by separately normalizing N7-G reactivities in 5’-GR and 5’-GY contexts. Final normalized protection scores *N_N7,i_* are computed as

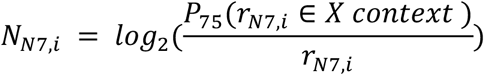

where *P*_75_(*r_N7,i_* ∈ *X context*) is the 75^th^ percentile of the N7-G reactivity distribution for either 5’-GR or 5’-GY contexts (matched to the sequence context of nucleotide *i*). For nucleotides with *r_N7,i_* < 0, indicating that the N7-G mutation rate is lower in the DMS-treated sample compared to unmodified sample, *N_N7,i_* is set to 3.32.

To facilitate robust normalization of short RNAs with <15 guanines in 5’-GR or <15 guanines in 5’-GY contexts, reactivities were heuristically adjusted and pooled together for calculation of the 75^th^ percentile normalization factors. Specifically, 5’-GR reactivities were multiplied by 1/0.65 and pooled with 5’-GY reactivities. For 5’-GY contexts, the 75^th^ percentile of this pooled distribution (*P*_75_(*pool*)) is used as the normalization factor. For 5’-GR contexts, 0.65\**P*_75_(*pool*) is used as the normalization factor.

As an automated quality control step, ShapeMapper requires the calculated N7-G normalization factor to be >0.015 for both 5’-GY and 5’-GR contexts to perform N7-G reactivity analysis.

All msDMS-MaP datasets were processed using ShapeMapper2.3 using the --min-mut-separation 0 --dms --N7 and --output-parsed-mutations flags. The --amplicon flag was used for amplicon libraries and the --random-primer-len 9 flag was used for randomly primed rRNA libraries. For the short RNAs TPP, U2, and P4-P6, the flag –pernt-norm-factor-threshold 15 was used to reduce the number of required guanines required for data normalization to 15 (the default requirement is 20 guanines).

For 7SK libraries labeled with UMIs, fastq files were pre-processed to remove duplicate reads prior to ShapeMapper processing. UMIs were extracted via UMI-tools^88^. The reads were then aligned to a reference fasta with Bowtie2^89^ using the parameters --local --sensitive-local --mp 3,1 --rdg 5,1 --dpad 30 --maxins 800 --ignore-quals --no-unal. The alignment file was sorted and indexed with UMI-tools, deduplicated using UMICollapse^90^, and then back-converted to paired fastq file format via UMI-tools. A script automating this deduplication pipeline can be downloaded at https://github.com/MustoeLab/umi_dedup_pipeline.

### ROC analysis

Area under the receiver operating characteristic curves (AUROC) for base-pairing status in S2 were calculated as previously described^18^. AUROC curves shown in Figure 2 were calculated with *Scikit-Learn* (0.24.1) in Python using normalized N7-G mutation rates where solvent accessible surface area (SASA) was obtained from the solved structure (PDB 7K00^85^). SASA was measured in PyMOL using a probe size of 4 Å. Solvent inaccessible N7-G positions were defined as positions with SASA < 1 Å^3^.

### Identification of *E. coli* rRNA self-folding tertiary domains

Putative self-folding tertiary domains were identified in protein-free, Mg^2+^-folded *E.coli* rRNA datasets by searching for sites featuring ≥4 N7-G protections (average protection score ≥1.6) within a contact distance^91^ of 10.

### PPV and sensitivity calculations

Positive predictive value (PPV) and sensitivity (sens) of N7-G protections were calculated in reference to N7-G solvent accessible surface area (SASA) measured in high-resolution molecular structures. Reference structures used were as follows: TPP (PDB 2GDI^32^), P4-P6 (PDB 1GID^92^), RNase P (PDB 6AHR^47^), U1 snRNA (PDB 6QX9^93^), tmRNA (PDB 7ABZ^42^), *E. coli* rRNA (PDB 7K00^85^). SASA was measured in PyMOL using a probe size of 4 Å. Solvent inaccessible N7-G positions were defined as positions with SASA < 1 Å^3^. Protected N7-G sites were defined as sites with average normalized protection score ≥1.6. Sites that were both solvent inaccessible and N7-G protected were defined as true positives (TP). Sites that were solvent accessible but N7-G protected were defined as false positives (FP). Finally, sites that were solvent inaccessible but not N7-G protected were defined as false negatives (FN). PPV and Sens were calculated as:

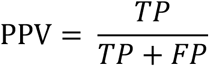

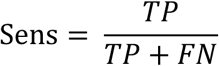

### RING calculation

We previously developed the RingMapper software suite for computing N1/3-N1/3 RINGs^30^. We updated RingMapper to permit concurrent calculation of N1/3-N1/3, N1/3-N7, and N7-N7 RINGs. RingMapper takes as input parsed N1/N3 and N7 mutation files output by ShapeMapper 2.3, which condense each sequencing read into binary vectors indicating whether each nucleotide position was measured, and if so, whether it was mutated in the N1/3 or N7 channels, respectively. Pairs of nucleotide positions that are jointly modified in a statistically significant correlated manner in the N1/3 channel, in the N1/3 channel for one nucleotide and the N7 channel for the other nucleotide, or in the N7 channel for both nucleotides are classified as N1/3-N1/3, N1/3-N7, and N7-N7 RINGs, respectively.

Prior versions of RingMapper quantified RINGs using the average product corrected G-statistic (G_APC_). However, G_APC_ is an imperfect metric because it scales with sequencing depth, meaning that it cannot be easily compared across samples sequenced to varied depths. Additionally, for deeply sequenced samples, even marginally correlated nucleotides can demonstrate significant G_APC_ scores, making it difficult to distinguish particularly strong RINGs. In this paper, we thus use a new metric, *c_ij_*, to quantify the coupling strength between nucleotides:

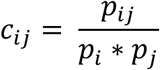

*p_ij_* is the joint probability of both nucleotides *i* and *j* being mutated in the considered N1/3-N1/3, N1/3-N7, or N7-N7 channels, and *p_i_* and *p_j_* are the marginal mutation probabilities of *i* and *j*. *c_ij_* reports the increase in co-mutations compared to random expectations, with *c_ij_*=2 denoting a two-fold enhanced rate of co-mutations. RINGs are reported between nucleotides if they pass statistical significance (G_APC_>20) and *c_ij_*>2. Analysis of multiple datasets revealed *c_ij_* to be reproducible across independent experiments and relatively invariant to read-depth.

### DANCE analysis

We updated the DANCE ensemble deconvolution software to permit concurrent determination of N1/3 and N7 reactivity profiles for co-existing RNA structural states in complex structural ensembles. Briefly, DANCE obtains both the reactivity profiles and populations of structural sub-states by fitting single-molecule DMS-MaP sequencing data to a modified Bernoulli mixture model ^52^. Fitting is performed for sequentially larger numbers of model states until the best fit is identified or until the maximum number of user-specified states is reached. Model fitting is performed in two stages. In the first stage, the mixture model is fit using a subset of “active” nucleotide positions that have high DMS signal-to-noise. In the second stage, reactivities for the remaining “inactive” nucleotide positions are solved using the “active”-fit mixture model as a constraint. We incorporated N7-G data into DANCE as additional “inactive” positions, thus effectively using deconvoluted N1/3 data as constraints to subsequently fit N7-G data. We also explored incorporating N7-G data as “active” during mixture model fitting but observed negligible benefits compared to the “inactive” strategy, accompanied by negative impacts of occasional reduced model fitting robustness and longer compute times. Following model fitting, raw N7-G reactivities are transformed into normalized protection scores following the strategy described above, with normalization factors computed using the maximum N7-G reactivity rates across all ensemble states. The updated version of the DanceMapper pipeline can be downloaded at https://github.com/MustoeLab/DanceMapper. For 7SK deconvolution was limited to a maximum of 3 states (--maxc 3) and for TPP, P4-P6 and U2 snRNA clustering was limited to 2 states (--maxc 2).

### 7SK structure modeling

For structure pairing probability calculations, data from three replicate experiments were combined to obtain single consolidated in-cell and protein-free datasets. Deduplicated reads were combined and then processed by ShapeMapper and DanceMapper. To increase power for detecting PAIRs, DanceMapper was run using the flag --readprob_cut 0.6. DMS restraints and PAIR data were then used as input for structural modeling with RNAstructure (v6.2) using the automated foldClusters.py script (distributed in the DanceMapper package). To prevent formation of erroneous long-range predicted pairs involving the terminal 3’ end, we constrained the terminal UUCUUUU nucleotides to be single-stranded during structural modeling (accomplished by coding these nucleotides as lowercase in the reactivities.txt file input into foldClusters.py).

### Data visualization with ArcPlot

N1/3 and N7 reactivity data, RINGs, and secondary structure arc diagrams were visualized using the ArcPlot software suite (https://github.com/MustoeLab/StructureAnalysisTools). ArcPlot was updated to permit visualization of N7-G protection scores and N1/3-N7 and N7-N7 RINGs. We also added new functionality to automate calculation and plotting of reactivity means and standard deviations from multiple experimental replicates.

### Δreactivity analysis

ΔN1/3 and ΔN7 reactivity analyses shown in Figure 6 and S9 were calculated as

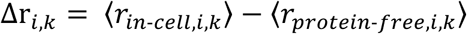

where Δr*_k,i_* is the reactivity difference of nucleotide *i* in DANCE-deconvoluted state *k,* and 〈*r_in-cell,i,k_*〉 and 〈*r_protein-free,i,k_*〉 denote the average normalized N1/3 or N7 reactivities of nucleotide *i* in state *k* measured in cells and protein-free experiments, respectively. Standard deviations, σ_Δ*r*_*i,k*__ were computed from error propagation as:

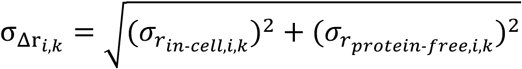

where *σ_rin-cell,i,k_* is the standard deviation of the normalized N1/3 or N7 reactivities across set of replicates. A two-sample T-test was used to identify nucleotides with significantly different reactivities across in-cell and cell-free experiments. Calculations used three independent in-cell msDMS-MaP replicates and three independent cell-free replicates.

### RNP-MaP experiments

RNP-MaP experiments were performed in replicates on separate days largely as described^14^. HEK293T cells were cultured in DMEM with 10% FBS, 100 U/ml penicillin, and 100 µg/ml streptomycin and grown to 80–90% confluency in 10-cm dishes. Cells were washed twice with PBS followed by addition of 4.75 ml of PBS and 250 µl of 200 mM SDA (NHS-diazirine, succinimidyl 4,4′-azipentanoate, Thermo Fisher) in DMSO or 250 µl of neat DMSO. Plates were incubated at 37 °C in 5% CO_2_ for 10 min for SDA labeling of proteins. PBS was then aspirated, semi-dry cells placed on ice, and proteins cross-linked to RNA by exposure to 3 J/cm^2^ of 365-nm-wavelength UV light (about 8 min in a crosslinker equipped with five 8-W F8T5 black lights) at a distance of ∼1 inch from the light source.

UV-treated (SDA-treated and untreated) cells were resuspended in 2.5 mL proteinase K lysis buffer (40 mM Tris-HCl (pH 8.0), 200 mM NaCl, 20 mM EDTA, 1.5% SDS and 0.5 mg/ml of proteinase K) and incubated 37 °C for 2 hr with intermittent mixing. RNA was recovered through two extractions with 25:24:1 phenol:chloroform:isoamyl alcohol and two extractions with chloroform and purified by isopropanol precipitation. Pellets were resuspended in 100 µl of 1× DNase buffer and incubated with 2 units of DNase (TURBO, Thermo Fisher Scientific) at 37 °C for 1 h. After the first incubation, 2 more units of TURBO DNase were added and samples incubated an additional 1 hr at 37 °C. RNA was purified using SPRI beads (Mag-Bind TotalPure NGS beads; Omega Bio-tek).

500 ng of total RNA RNA from SDA-treated or DMSO-treated cells was reversed transcribed using SuperScript II (Thermo Fisher) MaP conditions using 2 pmol 7SK-specific primer^14^. RT products were purified using columns (G-50 Microspin columns; Cytiva). 7SK sequencing libraries were then generated using a two-step PCR approach ^87^. 5 µl RT product was input as template for PCR1 (98 °C for 30 s; 20 cycles of 98 °C for 10 s, 66 °C for 30 s, 72 °C for 30 s; and 72 °C for 2 min; Q5 Hot Start DNA polymerase [NEB]). PCR1 product was purified by SPRI beads (Mag-Bind TotalPure NGS, Omega Bio-tek,; 1× ratio) and 2 ng of product was used as template for PCR2 (98 °C for 30 s; 10 cycles of 98 °C for 10 s, 66 °C for 30 s, 72 °C for 30 s; 2 min at 72 °C). PCR2 products were purified with SPRI beads (0.8× ratio) and sequenced on an Illumina MiSeq instrument using 2 × 250 paired-end sequencing (v2 chemistry).

RNP-MaP sequencing data were processed using ShapeMapper2.2 as described^14^. Primer binding regions and the 5 nucleotides immediately adjacent to the reverse primer binding site were not considered for analysis. Reactivity data represent the ratio of the mutation rates measured in SDA+UV and UV only samples, averaged over two replicates. Significant RNP-MaP sites were required to pass previously described nucleotide-specific reactivity thresholds and z-factor cutoffs in both independent replicates^14^.

