## Supplementary Information for "Simultaneous mapping of RNA secondary, tertiary, and quaternary structure in living cells by multi-site DMS probing"

Supporting figures (10), Supporting table (1)

#### Figure S1

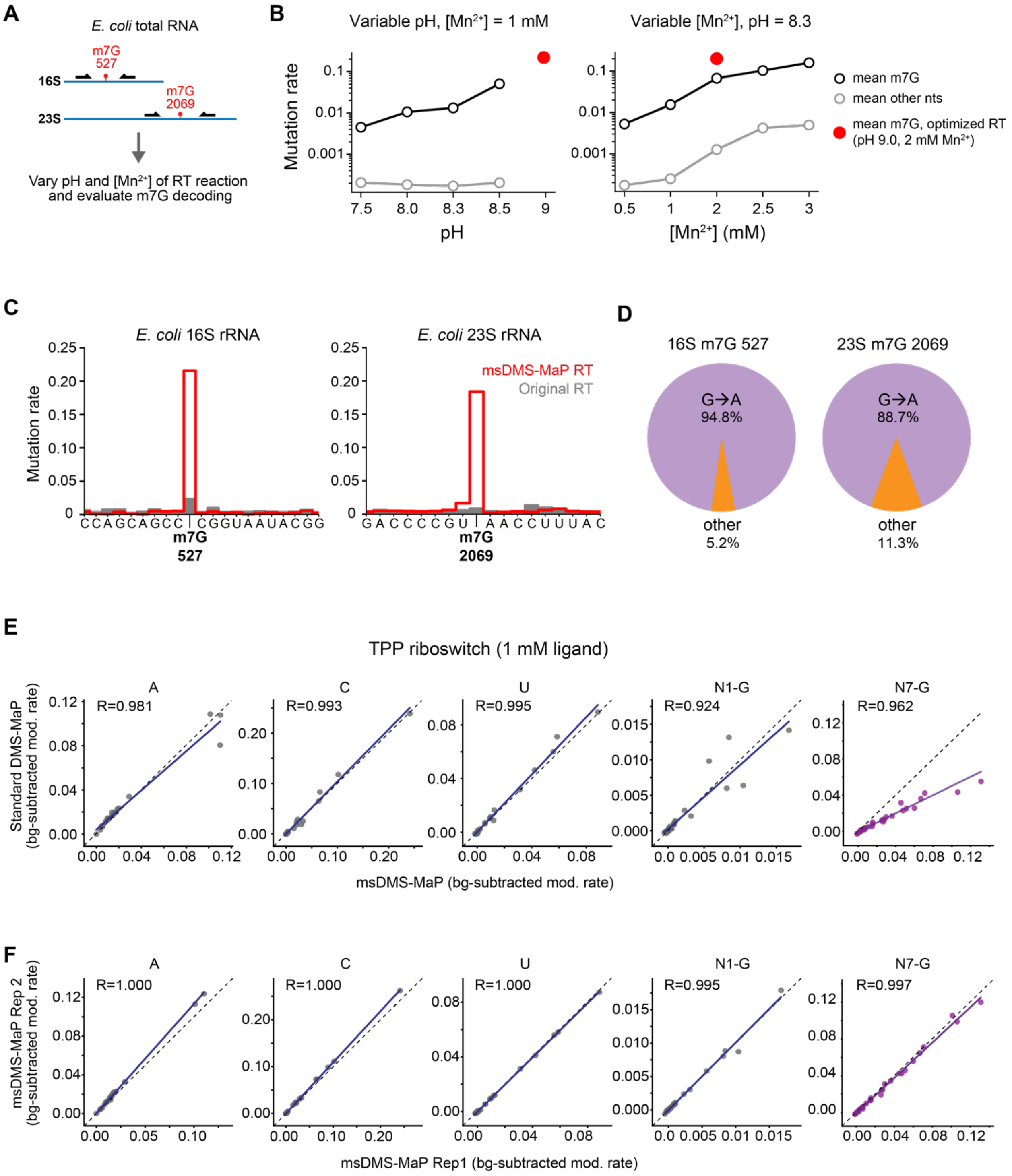

**Figure S1: N7-G decoding efficiency depends on pH and $Mn^{2+}$ concentration during reverse transcription.** Changes to the RT reaction produce minor changes in background reactivity rates and are highly reproducible. (A) Schematic of the library design used to evaluate m7G decoding. (B) Comparison of mutation rates at natural m7G positions in unmodified *E. coli* rRNA with variable pH at a fixed manganese concentration (left) and variable manganese concentration and fixed pH (right). The final selected condition is shown in red. (C) Comparison of mutation rates in unmodified *E. coli* rRNA using msDMS-MaP RT and standard SSII MaP RT. (D) Breakdown of m7G decoding in unmodified *E.coli* rRNA. (E) Comparison of standard Marathon DMS-MaP versus msDMS-MaP reactivities at all four bases and the N7-G position in the TPP riboswitch. Pearson's R is shown. (F) Comparison of TPP riboswitch msDMS-MaP reactivities in two replicates. Pearson's R is shown.

#### Figure S2

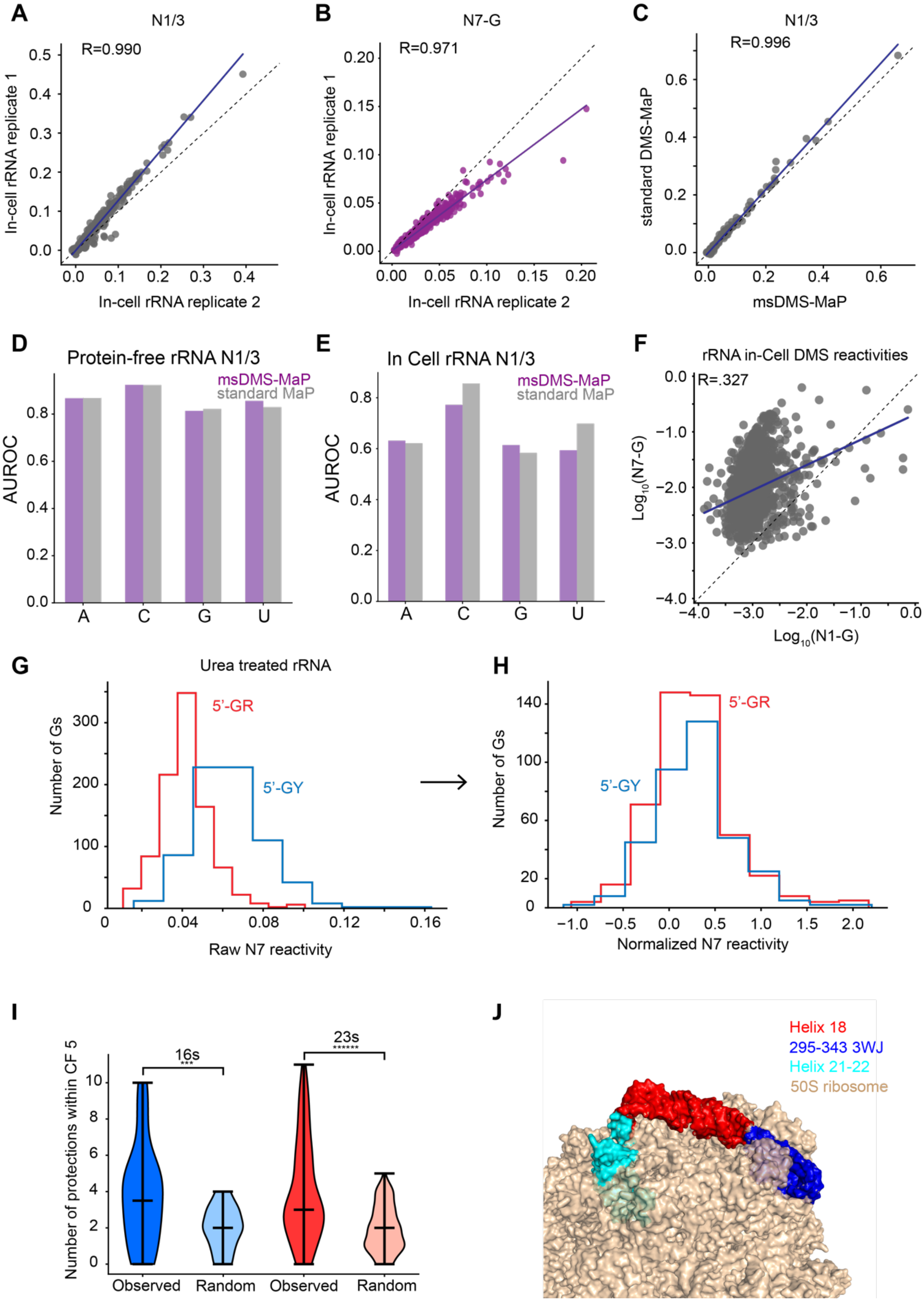

A
N1/3
R=0.990
In-cell rRNA replicate 1
In-cell rRNA replicate 2
B
N7-G
R=0.971
In-cell rRNA replicate 1
In-cell rRNA replicate 2
C
N1/3
R=0.996
standard DMS-MaP
msDMS-MaP
D
Protein-free rRNA N1/3
msDMS-MaP
standard MaP
AUROC
A
C
G
U
E
In Cell rRNA N1/3
msDMS-MaP
standard MaP
AUROC
A
C
G
U
F
rRNA in-Cell DMS reactivities
R=.327
$Log_{10}$(N7-G)
$Log_{10}$(N1-G)
G
Urea treated rRNA
5'-GR
5'-GY
Number of Gs
Raw N7 reactivity
H
5'-GR
5'-GY
Number of Gs
Normalized N7 reactivity
I
16s
23s
Number of protections within CF 5
Observed
Random
Observed
Random
J
Helix 18
295-343 3WJ
Helix 21-22
50S ribosome

**Figure S2: msDMS-MaP provides orthogonal measurements that inform on complex RNA structure while maintaining the ability to measure RNA secondary structure.** (A) Comparison of N1/3 and (B) N7 *E. coli* rRNA reactivities using msDMS-MaP in two replicates. Pearson's R is shown. (C) Comparison of N1/3 *E. coli* rRNA reactivities using standard MarathonRT at pH 8.3 versus msDMS-MaP. Pearson's R is shown. (D, E) Computed AUROC statistics for N1/3 reactivities ability to distinguish between paired and unpaired nucleotides at each base in (D) protein-free and (E) in-cell *E. coli* rRNA. msDMS-MaP is shown in purple and standard SSII MaP is shown in gray. (F) Measured N1-G reactivity versus N7-G reactivity of *E. coli* rRNA guanines. Pearson's R is shown. Distribution of raw (G) and normalized (H) N7-G reactivity rates based on the identity of 3' neighboring nucleotide. (I) Clustering of N7-G protections within close secondary structure proximity (contact distance $\leq$5) measured in protein-free rRNA versus if protections are randomly shuffled across rRNA guanines. Significance was computed using a two-sided Mann-Whitney U test; **, $p<10^{-3}$, ******, $p<10^{-6}$. (J) *E. coli* ribosome structure highlighting helix 18 in red, 295-343 3WJ in dark blue, and helices 21 and 22 in cyan relative to the rest of the 50S ribosome in brown.

#### Figure S3

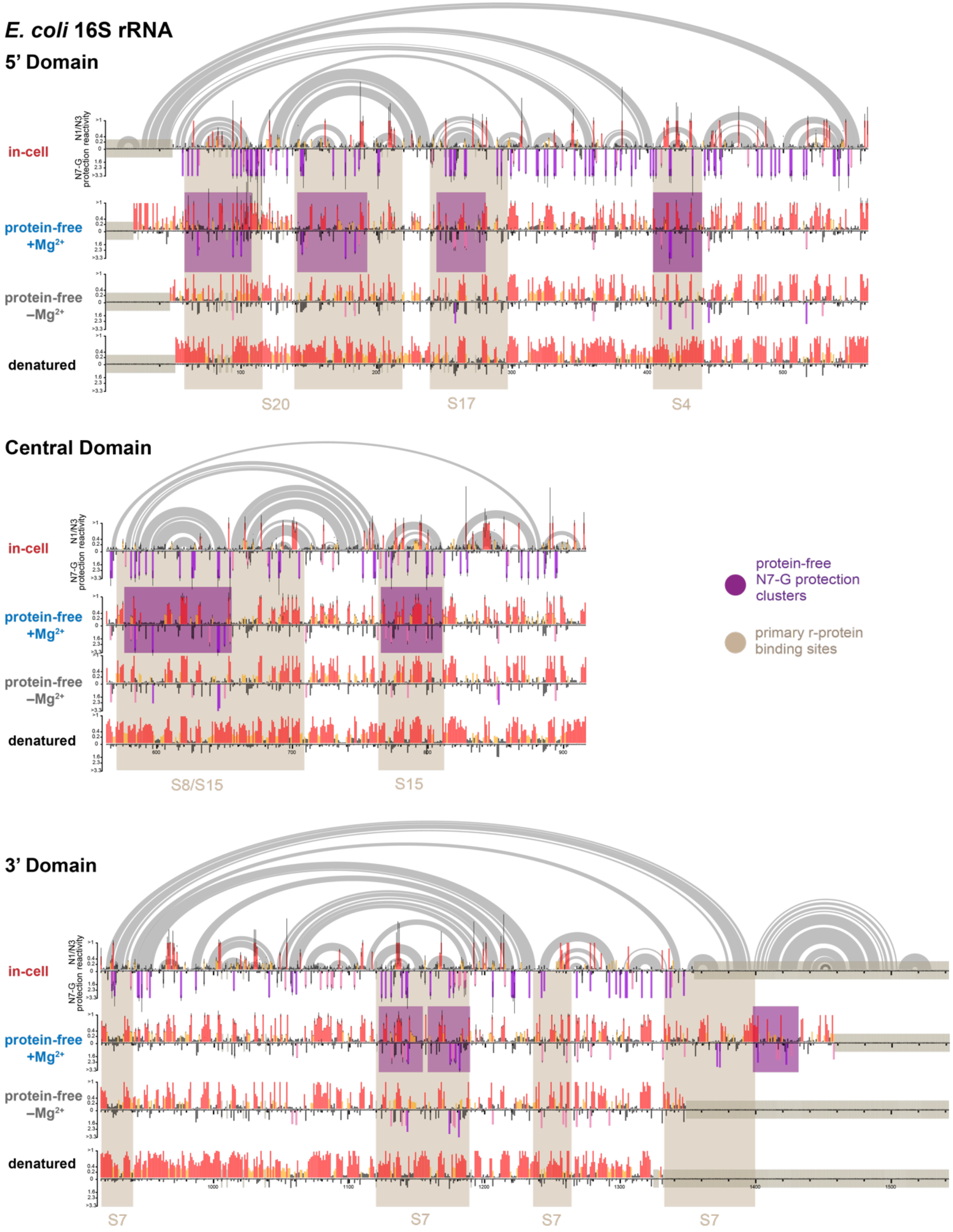

**Figure S3: msDMS-MaP reactivity profiles for 16S rRNA under varied folding conditions.** Self-folding domains identified based on clustered N7-G protections in protein-free +$Mg^{2+}$ are highlighted in purple. Primary r-protein binding sites are highlighted in brown.

#### Figure S4 — part I

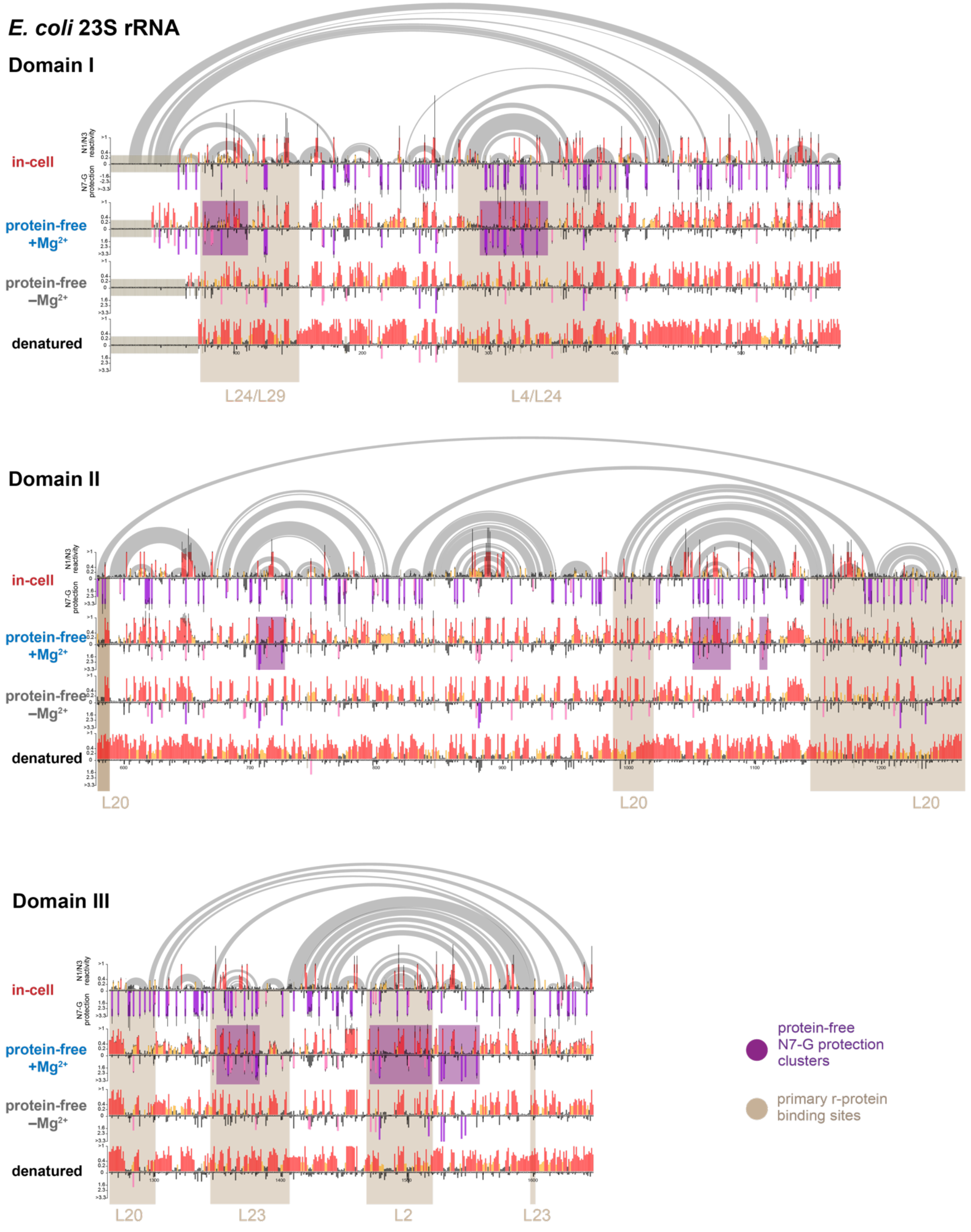

### Figure S4 — part II

#### *E. coli* 23S rRNA

##### Domain IV

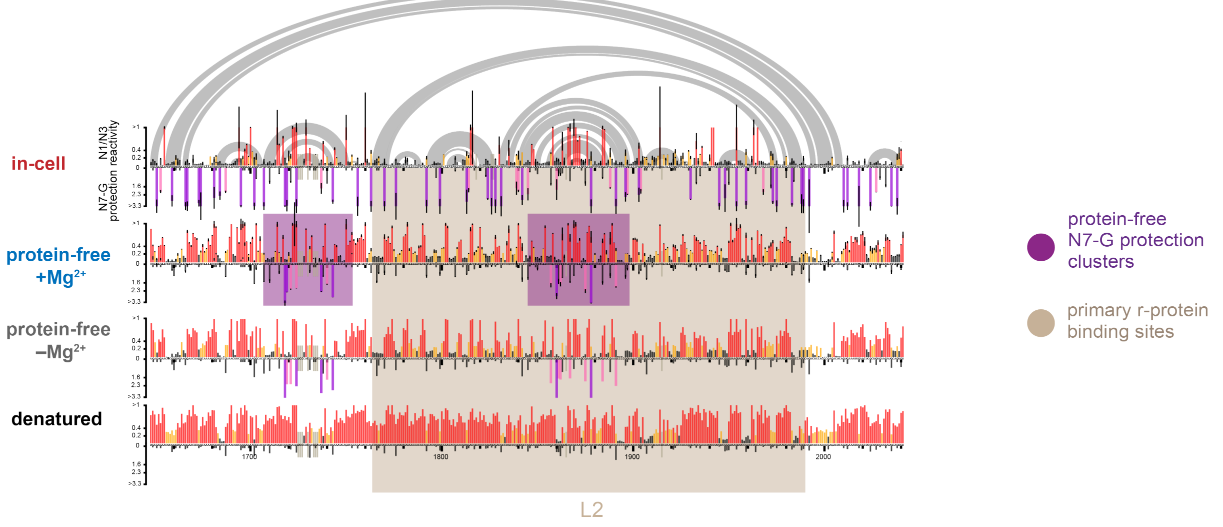

##### Domain V

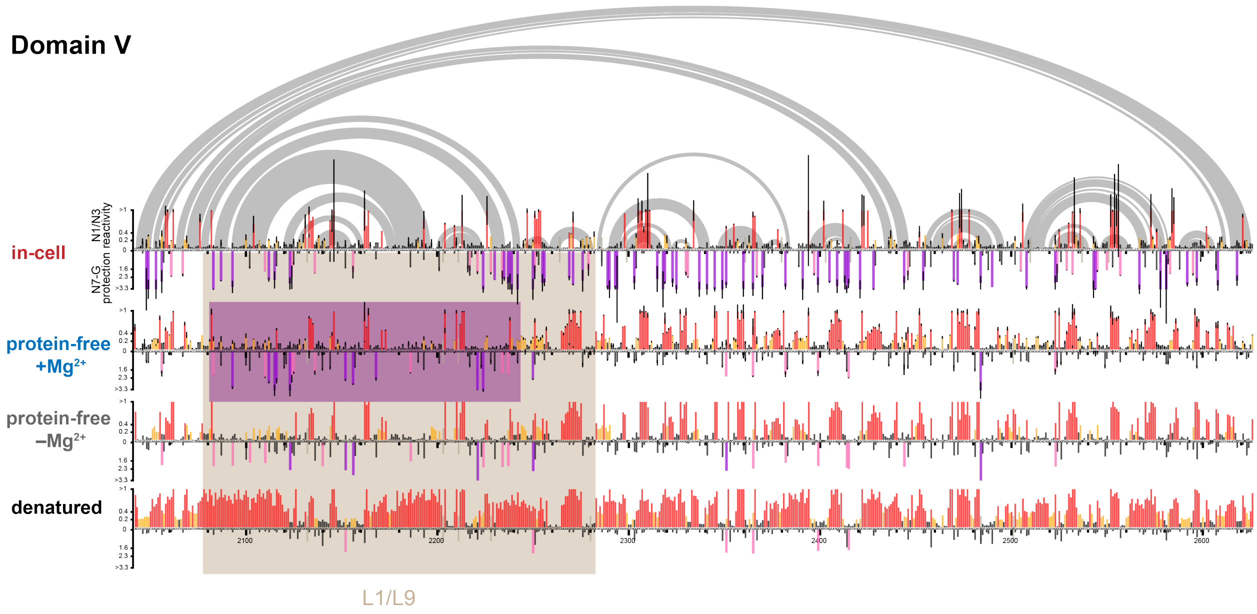

##### Domain VI

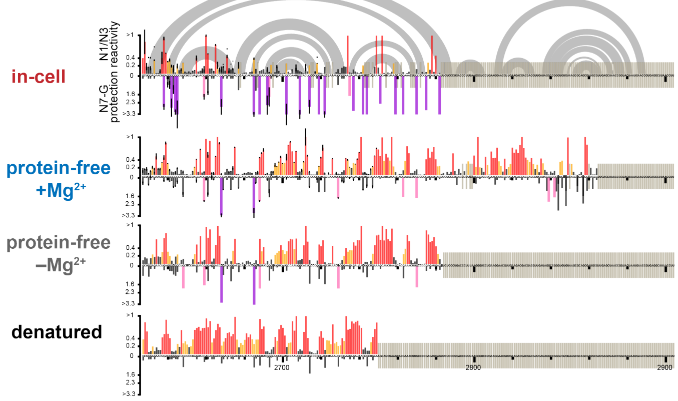

**Figure S4: msDMS-MaP reactivity profiles for 23S rRNA under varied folding conditions.** Self-folding domains identified based on clustered N7-G protections in protein-free +$Mg^{2+}$ are highlighted in purple. Primary r-protein binding sites are highlighted in brown.

#### Figure S5

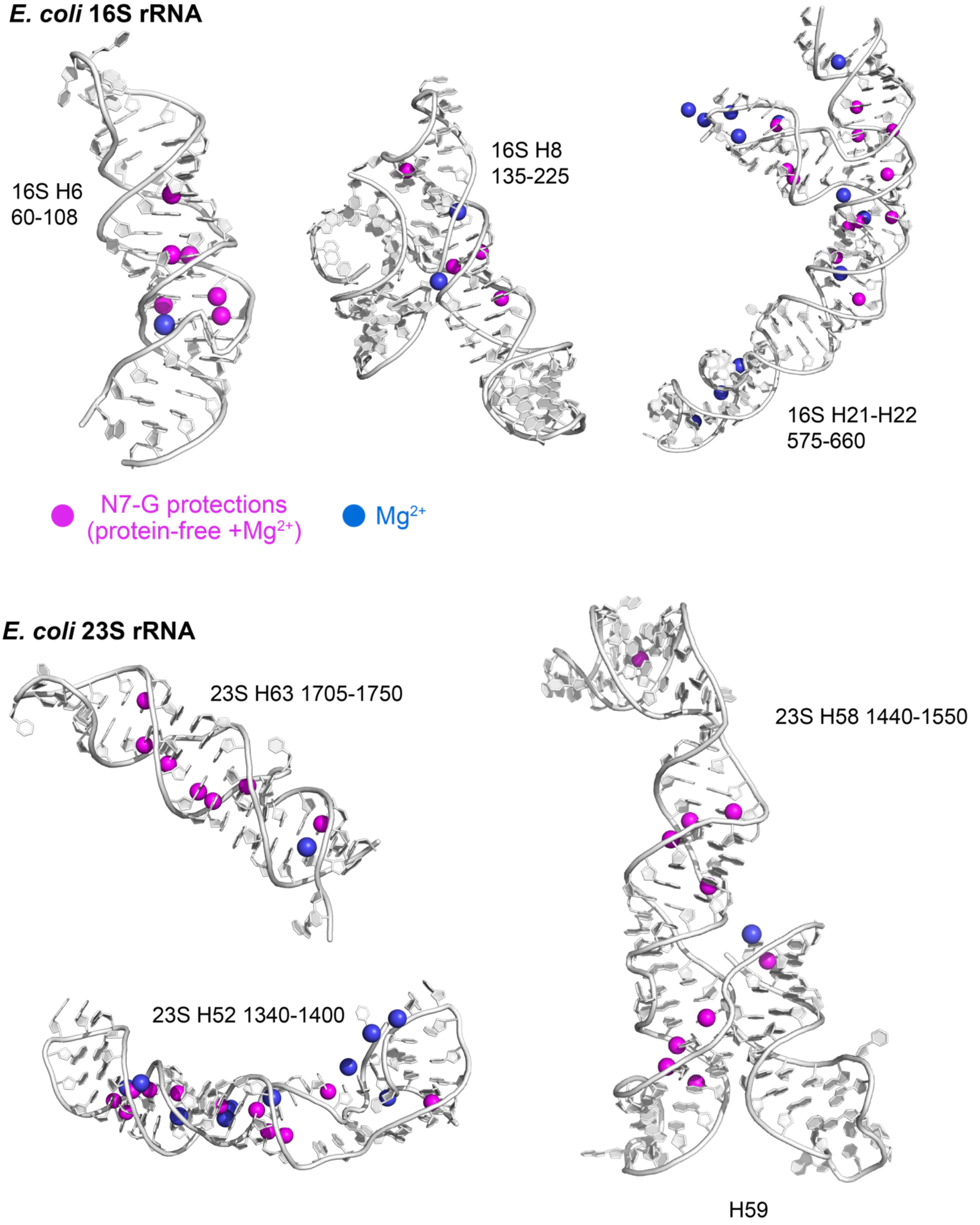

**Figure S5: Example *E. coli* rRNA independent folding domains identified by msDMS-MaP.** Structures are excerpted from PDB 7K00. N7-G protections measured in protein-free +$Mg^{2+}$ \rRNA are highlighted in purple. Magnesium ions are shown in blue.

#### Figure S6

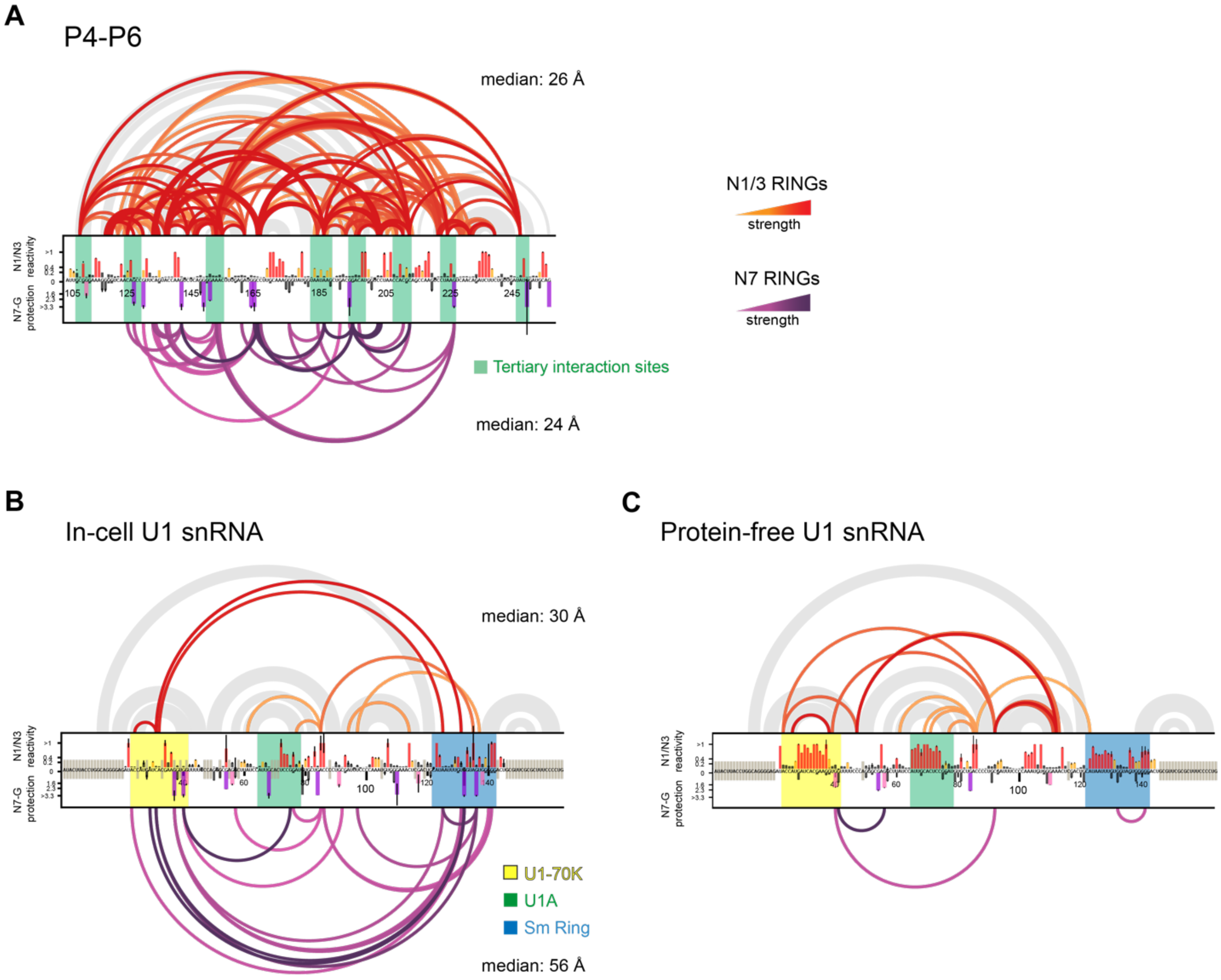

**Figure S6: RINGs identify coupled tertiary folding and protein binding sites.** Correlated N1/3 and N7 mutations in (A) P4-P6 domain of the *Tetrahymena* ribozyme with $Mg^{2+}$, (B) in-cell human U1 snRNA, and (C) protein-free human U1 snRNA. RINGs were required to be observed in two independent replicates. N1/3 and N7-G reactivity profiles represent the average of two replicates, with error bars denoting the standard deviation. Protein binding sites and tertiary interaction sites are highlighted. Gray arcs at top indicate base pairing interactions. Darker colors indicate stronger RINGs. For systems with a solved structure (A and B), median three-dimensional distances spanned by N1/3 and N7 RINGs are listed at top and bottom, respectively.

#### Figure S7

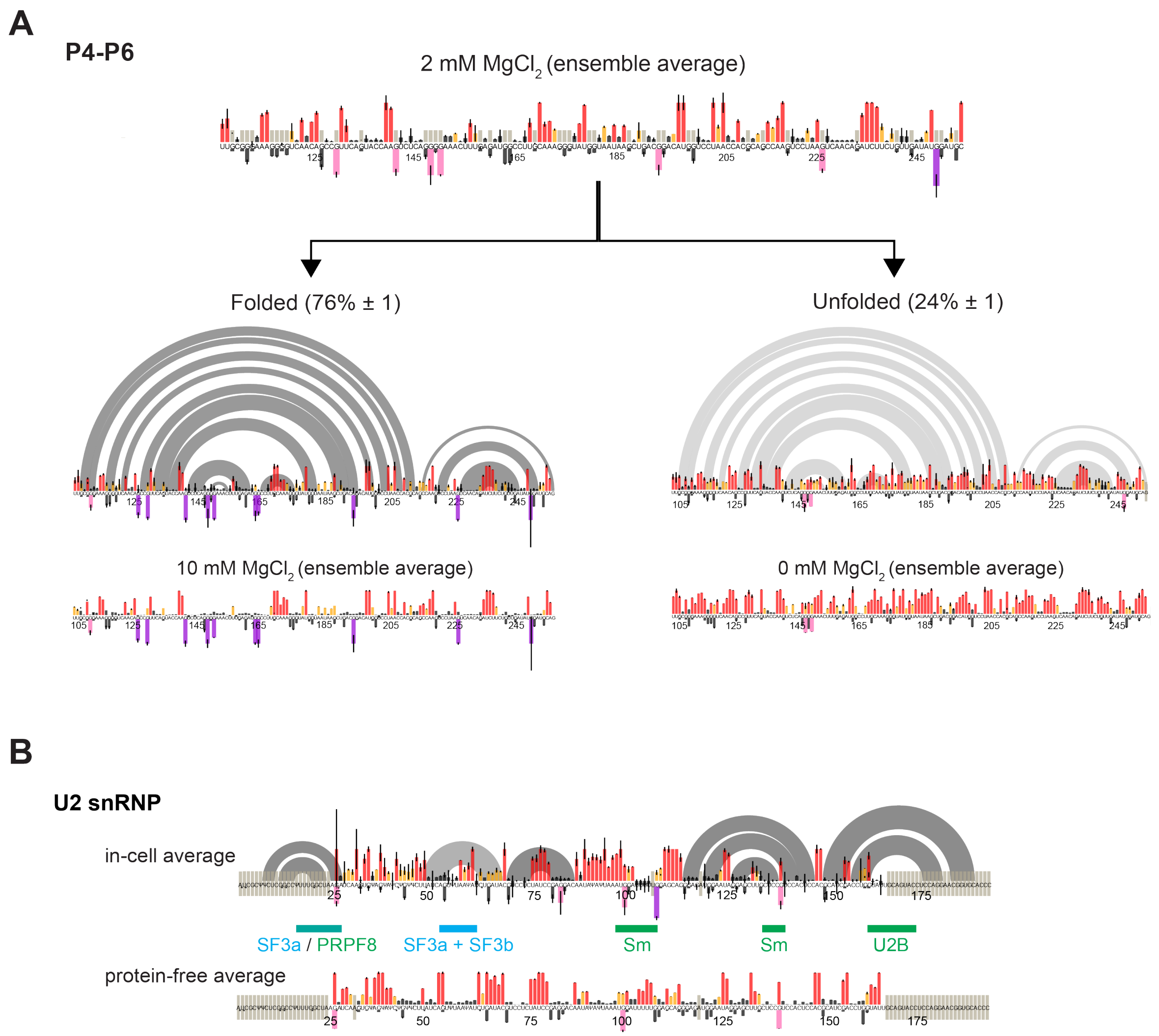

**Figure S7: Measurement of state-specific N7-G protections using msDMS-MaP and DANCE deconvolution.** (A) msDMS-MaP experiments were performed on the P4-P6 domain of the *Tetrahymena* ribozyme folded with sub-saturating 2 mM $Mg^{2+}$. DANCE resolves the RNA into tertiary folded and unfolded states with N1/3 and N7-G reactivity profiles that closely match the profiles observed when P4-P6 is probed under fully unfolded (0 mM $Mg^{2+}$) or fully folded (10 mM $Mg^{2+}$) conditions. Data represent averages over two independent replicates, with error intervals denoting the standard deviation. (B) Averaged (non-deconvoluted) msDMS-MaP reactivity profiles of human U2 snRNA from RPE-1 cells probed in cells and under protein-free conditions. Data represent averages over two independent replicates, with error bars denoting the standard deviation.

#### Figure S8

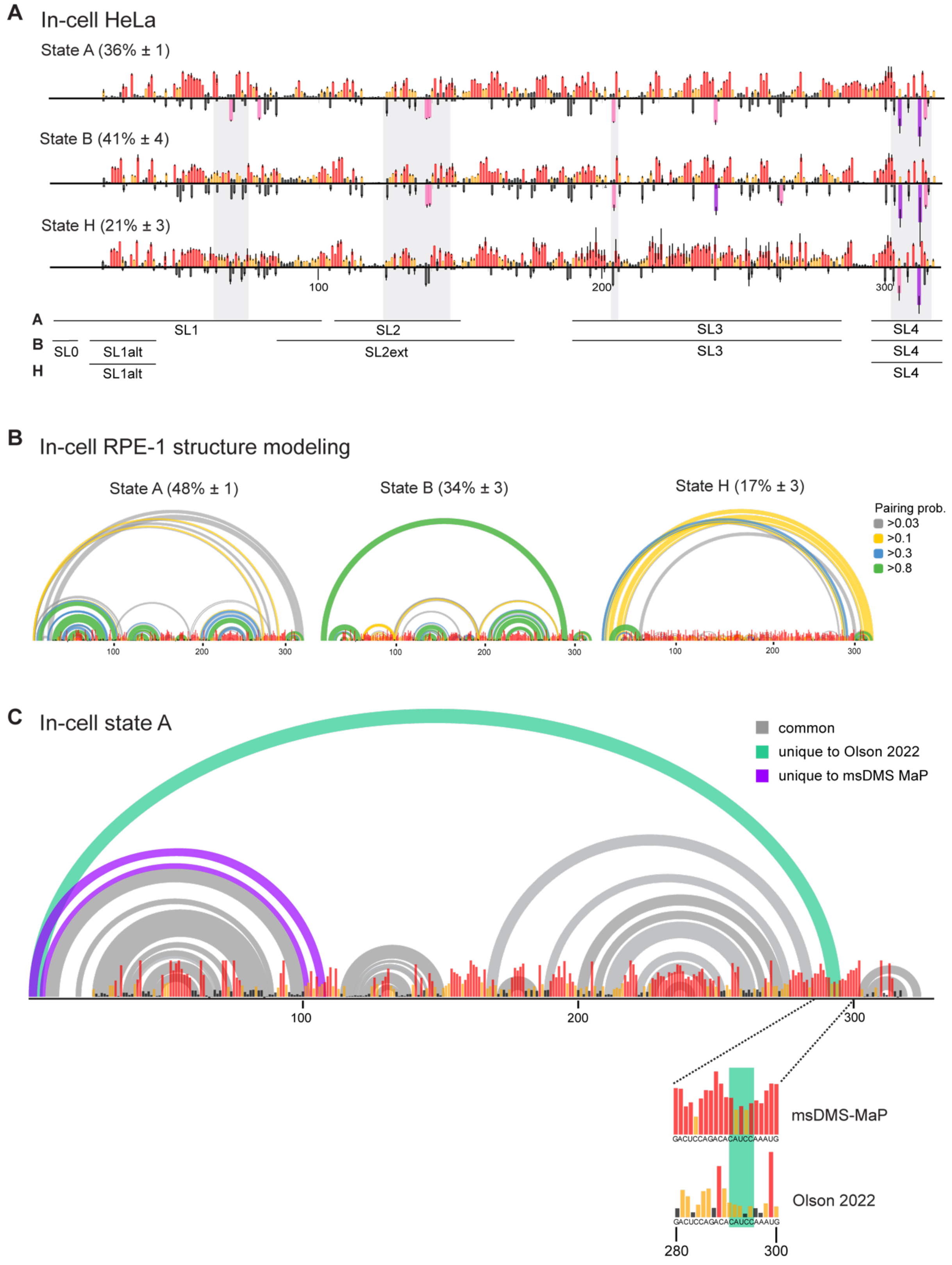

A
In-cell HeLa
State A (36% ± 1)
State B (41% ± 4)
State H (21% ± 3)
100
200
300
A
B
H
SL1
SL2
SL3
SL4
SL0
SL1alt
SL2ext
SL3
SL4
SL1alt
SL4
B
In-cell RPE-1 structure modeling
State A (48% ± 1)
State B (34% ± 3)
State H (17% ± 3)
Pairing prob.
>0.03
>0.1
>0.3
>0.8
100
200
300
C
In-cell state A
common
unique to Olson 2022
unique to msDMS MaP
100
200
300
msDMS-MaP
Olson 2022
GACUCCAGACACAUCCAAAUG
280
300

**Figure S8: 7SK structural ensemble measured in HeLa and RPE-1 cells.** (A) DANCE-deconvoluted N1/3 and N7-G reactivity profiles of 7SK RNA measured in HeLa cells. Data represent averages over two replicates, with error bars denoting standard deviations. Nucleotides protected in RPE-1 cells (Figure 6) are highlighted in grey. (B) Structural models for states A, B, and H guided by RPE-1 in-cell msDMS-MaP data. Displayed reactivity data, which was used to guide structure modeling, was obtained by combining three independent in-cell replicates. Modeled base-pairing probabilities are shown at top as arcs. (C) Comparison of current revised consensus model of state A with previously published consensus model[52]. Shown inset is a comparison between current msDMS-MaP data and prior DMS-MaP data in the SL0 forming region. Prior 7SK datasets were collected using SSII RT, which has reduced accuracy[18] compared to MarathonRT used for msDMS-MaP.

#### Figure S9

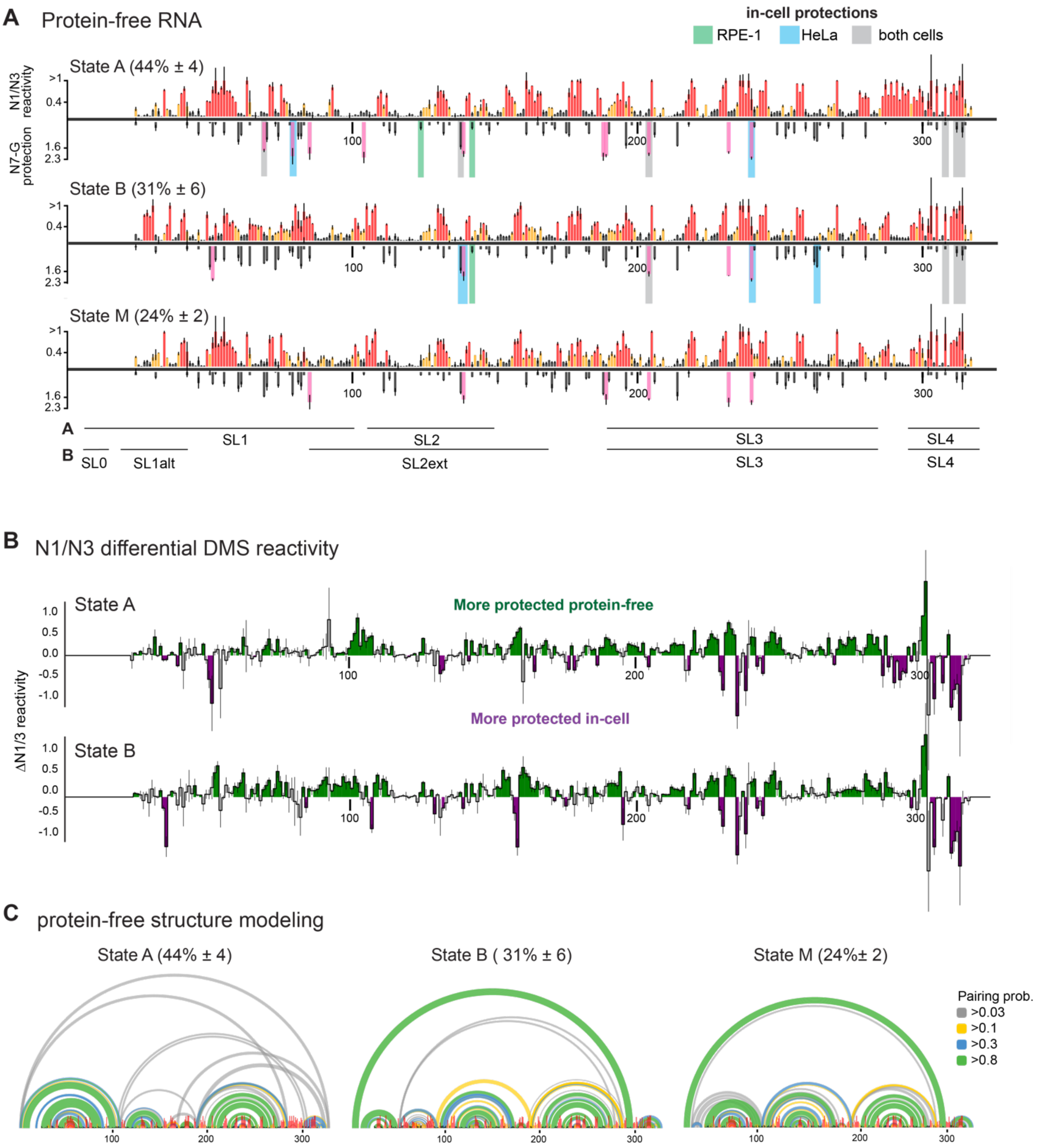

**Figure S9: 7SK structural ensemble under protein-free conditions.** (A) DANCE-deconvoluted N1/3 and N7-G reactivity profiles of 7SK RNA measured under protein-free conditions. Data represent averages over two replicates, with error intervals denoting standard deviations. Sites of N7-G protection observed in RPE-1 cells and HeLa cells are highlighted. (B) msDMS-MaP-supported structural models for protein-free states A, B, and M. Displayed reactivity data, which was used to guide structure modeling, was obtained by combining three independent protein-free replicates. Modeled base-pairing probabilities are shown at top as arcs. (C) Differential analysis of N1/3 reactivity in cells versus protein-free 7SK, showing State A (top) and State B (bottom). Differences represent averages between three in-cell and three protein-free replicates, with error bars denoting standard deviations. Nucleotides exhibiting statistically significant ($p<0.05$) increases in N1/3 reactivity in cells are shown in purple, and statistically significant increases in protein-free conditions in green.

#### Figure S10

##### Submission script to ShapeMapper

```
/storage/mustoe/software/N7_Release/shapemapper2.3-release/shapemapper \
--nproc 8 --name Incell_U2_snRNA_REP1 --target U2_snRNA.fa --amplicon --dms --serial --N7 --min-mutation-separation 0 \
--output-parsed --overwrite --temp U2_REP1 --out U2_snRNA \
--modified --R1 Incell_U2_snRNA_Mod_S3_L001_R1_001.fastq --R2 Incell_U2_snRNA_Mod_S3_L001_R2_001.fastq \
--untreated --R1 Incell_U2_snRNA_Unmod_S3_L001_R1_001.fastq --R2 Incell_U2_snRNA_Unmod_S3_L001_R2_001.fastq
```

##### Output files from ShapeMapper

Incell_U2_snRNA_REP1.fa
Incell_U2_snRNA_REP1_Modified_U2_snRNA_parsed.mut
Incell_U2_snRNA_REP1_Modified_U2_snRNA_parsed.mutga
Incell_U2_snRNA_REP1_U2_snRNA.dms
Incell_U2_snRNA_REP1_U2_snRNA_histograms.pdf
Incell_U2_snRNA_REP1_U2_snRNA.map
Incell_U2_snRNA_REP1_U2_snRNA_mapped_depths.pdf
Incell_U2_snRNA_REP1_U2_snRNA_per-amplicon_abundance.txt
Incell_U2_snRNA_REP1_U2_snRNA_profiles.pdf
Incell_U2_snRNA_REP1_U2_snRNA_profile.txt
Incell_U2_snRNA_REP1_U2_snRNA_profile.txtga
Incell_U2_snRNA_REP1_U2_snRNA_ribosketch_colors.txt
Incell_U2_snRNA_REP1_U2_snRNA_varna_colors.txt
Incell_U2_snRNA_REP1_Untreated_U2_snRNA_parsed.mut
Incell_U2_snRNA_REP1_Untreated_U2_snRNA_parsed.mutga

**new features**

##### Reactivity profile

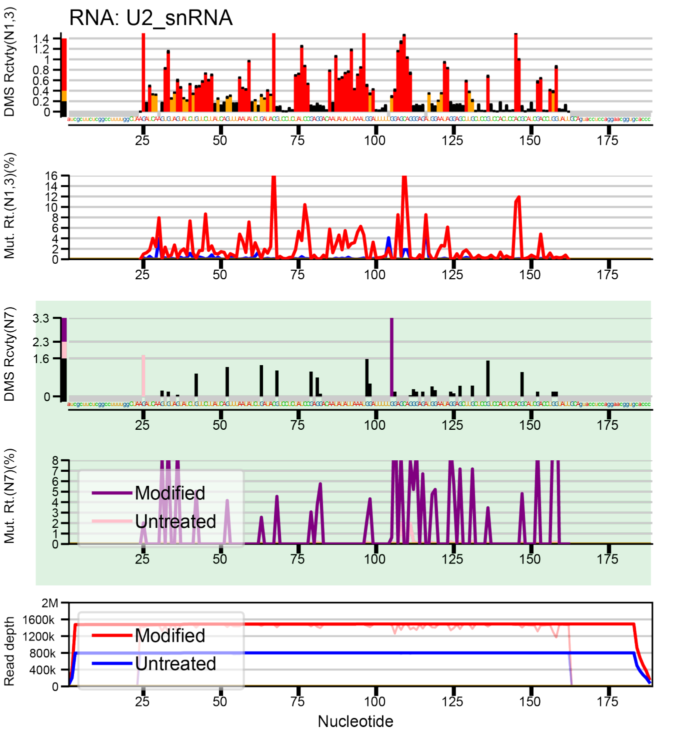

**Figure S10: Automated msDMS-MaP analysis using ShapeMapper2.3.** Example input and output files are shown, with newly added features highlighted in green.

**Table S1**: Primer and template sequences

| Name of primer | Sequence |
|---|---|
| 7SK_RT primer with LNA (+ indicates LNA base) | CCTACACGACGCTCTTCCGATCTTNNNNNNNNNNAA+GAAA+GG+CAGAC |
| 7SK_PCR1_Forward | GACTGGAGTTCAGACGTGTGCTCTTCCGATCTNNNNNGGATGTGAGGGCGATCTGG |
| 7SK_PCR1_Reverse | CCC TAC ACG ACG CTC TTC CGA TCT |
| TPP and P4-P6 invitro_RT primer | GGTTGTTTCTTTGGTTTGGTTTTG |
| invitro_PCR1_Forward | GACTGGAGTTCAGACGTGTGCTCTTCCGATCTNNNNNGGCCAAAACAACAACCGG |
| invitro_PCR1_Reverse | CCCTACACGACGCTCTTCCGATCTNNNNNGGTTGTTTCTTTGGTTTGGTTTTG |
| U1_RT_primer | CAGGGGAAAGCGCGAA |
| U1_PCR1_Forward | GACTGGAGTTCAGACGTGTGCTCTTCCGATCTNNNNNATACTTACCTGGCAGGG |
| U1_PCR1_Reverse | CCCTACACGACGCTCTTCCGATCTNNNNNCAGGGGAAAGCGCGAA |
| U2_RT_primer | GGGTGCACCGTTCCTGGAGGTAC |
| U2_PCR1_Forward | GACTGGAGTTCAGACGTGTGCTCTTCCGATCTNNNNNATCGCTTCTCGGCCTTTTGG |
| U2_PCR1_Reverse | CCCTACACGACGCTCTTCCGATCTNNNNNGGTGCACCGTTCCTGGAGGTAC |
| RNase P RT primer | AATGGGCGGAGGAGAGTAG |
| RNaseP_PCR1_Forward | CCCTACACGACGCTCTTCCGATCTNNNNNATAGGGCGGAGGGAAGC |
| RNaseP_ PCR1_Reverse | GACTGGAGTTCAGACGTGTGCTCTTCCGATCTNNNNNNAATGGGCGGAGGAGAGTAG |
| E. coli tmRNA _RT primer | GAGCTGGCGGGAGTTGAA |
| E. coli tmRNA_PCR1_Forward | CCCTACACGACGCTCTTCCGATCTNNNNNCTGGATTCGACGGGATTTGC |
| E. coli tmRNA_PCR1_Reverse | GACTGGAGTTCAGACGTGTGCTCTTCCGATCTNNNNNGAGCTGGCGGGAGTTGAA |
| E. coli Amplicon 23S_RT_primer | GCGTCCACACTTCAAAGCC |
| E. coli Amplicon 23S_PCR1_Forward | GACTGGAGTTCAGACGTGTGCTCTTCCGATCTNNNNNCACCCGAGACTCAGTGAAATTG |
| E. coli Amplicon 23S_PCR1_Reverse | CCCTACACGACGCTCTTCCGATCTNNNNNGCGTCCACACTTCAAAGCC |
| E. coli Amplicon 16S_RT primer | AGGTTGAGCCCGGGGATTTC |

| | |
|---|---|
| E. coli Amplicon 16S_PCR1_Forward | GACTGGAGTTCAGACGTGTGCTCTTCCGATCTNNN<br>NNCTCATTGACGTTACCCGCAGA |
| E. coli Amplicon 16S_PCR1_Reverse | CCCTACACGACGCTCTTCCGATCTNNNNNAGGTTG<br>AGCCCGGGGATTTC |
| **IVT templates** | |
| TPP_*invitro*_template with T7 binding site | TTCTAATACGACTCACTATAGGCCAAACAACAACC<br>GGGCCA AGGAC TCGGG GTGCC CTTCT GCGTG<br>AAGGC TGAGA AATAC CCGTA TCACC TGATC<br>TGGAT AATGC CAGCG TAGGG AAGTT CTCGA<br>TCCGG TTCGC CGGAT<br>CC AAAACCAAACCAAAGAAACAACC |
| P546_*invitro*_template with T7 binding site | TTCTAATACGACTCACTATAGGCCAAAACAACAACC<br>GGAATTGCGGGAAAGGGGTCAACAGCCGTTCAGTA<br>CCAAGTCTCAGGGGAAACTTTGAGATGGCCTTGCA<br>AAGGGTATGGTAATAAGCTGACGGACATGGTCCTA<br>ACCACGCAGCCAAGTCCTAAGTCAACAGATCTTCTG<br>TTGATATGGATGCAGTTCAAAACCAAACCAAAGAAA<br>CAACC |
